# Molecular Machines Stimulate Intercellular Calcium Waves and Cause Muscle Contraction

**DOI:** 10.1101/2022.11.04.515191

**Authors:** Jacob L. Beckham, Alexis R. van Venrooy, Soonyoung Kim, Gang Li, Bowen Li, Guillaume Duret, Dallin Arnold, Xuan Zhao, Ana L. Santos, Gautam Chaudhry, Jacob T. Robinson, James M. Tour

## Abstract

Intercellular calcium waves (ICW) are complex signaling phenomena that control many essential biological activities, including smooth muscle contraction, vesicle secretion, gene expression, and changes in neuronal excitability. Accordingly, the remote stimulation of ICW may result in versatile new biomodulation and therapeutic strategies. Here, we demonstrate that light-activated molecular machines (MM), molecules that rotate and perform mechanical work on the molecular scale, can remotely stimulate ICW. Live-cell calcium tracking and pharmacological experiments reveal that MM-induced ICW are driven by the activation of inositol triphosphate (IP3) mediated signaling pathways by unidirectional, fast-rotating MM. We then demonstrated that MM-induced ICW can be used to control muscle contraction *in vitro* in cardiomyocytes and animal behavior *in vivo* in *Hydra vulgaris*. Consequentially, this work demonstrates a new strategy for the direct control of cell signaling and downstream biological function using molecular-scale devices.

## Main

Calcium signaling impacts nearly every process relevant to cellular life, and the ability of calcium ions to alter protein shape and charge by reversible binding constitutes the most ubiquitous signaling motif in receptor biology^1^. The localized nature of calcium signaling, as well as its ability to activate downstream effector proteins, allows it to drive a vast array of biological activities. In single cells, calcium directly controls cellular proliferation^2^, gene expression^3,4^, differentiation^5^, movement^6^, and metabolism^7^. In organisms, calcium signals propagate through secondary messengers to cause intercellular calcium waves (ICW) that coordinate concerted action in whole tissues^8^. ICW play direct or indirect roles in processes ranging from muscle contraction^9^, potentiation of neuronal firing^10^, blood vessel dilation^11^, digestion^12^, and breathing^13^. Dysfunctional calcium signaling contributes to disease states ranging from cancer to cardiovascular disease and neurodegenerative disorders^14-15^.

Due to the multiplexed nature of calcium signaling^16^, the ability to remotely trigger ICW with high spatiotemporal precision may permit access to numerous downstream signaling pathways, offering a dynamic new strategy for the control of biological activity. Such advances may also yield new therapeutic avenues for diseases characterized by calcium signaling dysfunction. Currently, ICW are largely initiated experimentally by chemical methods^17^ or by applying mechanical stimuli using a micropipette attached to a microcontroller^2,8,18^.

Here, we describe the generation of ICW *via* the nanomechanical action of a light-activated molecular machine. Molecular machines (MM) are molecules that can be activated by external stimuli, such as light, to perform mechanical work on the molecular scale^19^. Just as the mechanical perturbation of a cell’s outer membrane causes intracellular calcium responses, the application of mechanical force *via* a fast, unidirectionally rotating MM elicits calcium release from the endoplasmic reticulum (ER). Slow-rotating and non-unidirectional MM do not elicit calcium flux under the same conditions, implicating a mechanism of action that depends on rapid, unidirectional molecular rotation. Calcium release is attenuated by the inhibition of inositol triphosphate (IP_3_)-mediated signaling, suggesting the IP_3_ signaling pathway as a primary driver of cellular responses. Finally, we show that MM-induced ICW can be exploited to control downstream biological processes, like muscle contraction, both *in vitro* and *in vivo*.

## Results and Discussion

### Fast, Unidirectional MM Induce Calcium Waves

MM employed in this study have overcrowded alkene motors based on the primary design by Feringa *et al*^19^. Their typical structure consists of a rotor connected to a stator by an atropisomeric alkene. When these MM are excited by incident photons, the rotor rotates unidirectionally relative to the stator, undergoing two photoisomerization steps and two thermal helix inversions before returning to the starting position (Fig. 1a). Overcrowded alkene motors locomote in solution^20^, drill through synthetic lipid bilayers and cell membranes^21^, and have previously been used to exert mechanical forces on individual cell-surface receptors *via* antibody targeting^22^. Here, we study cellular responses to MM administered without the use of chemical targeting or extracellular scaffolds, facilitating their use *in vivo*. The moiety “X” in Fig. 1a can be interchanged to modulate MM rotation rate, which is determined by the favorability of the thermal helix inversion step^23,24^.

**Fig. 1.**
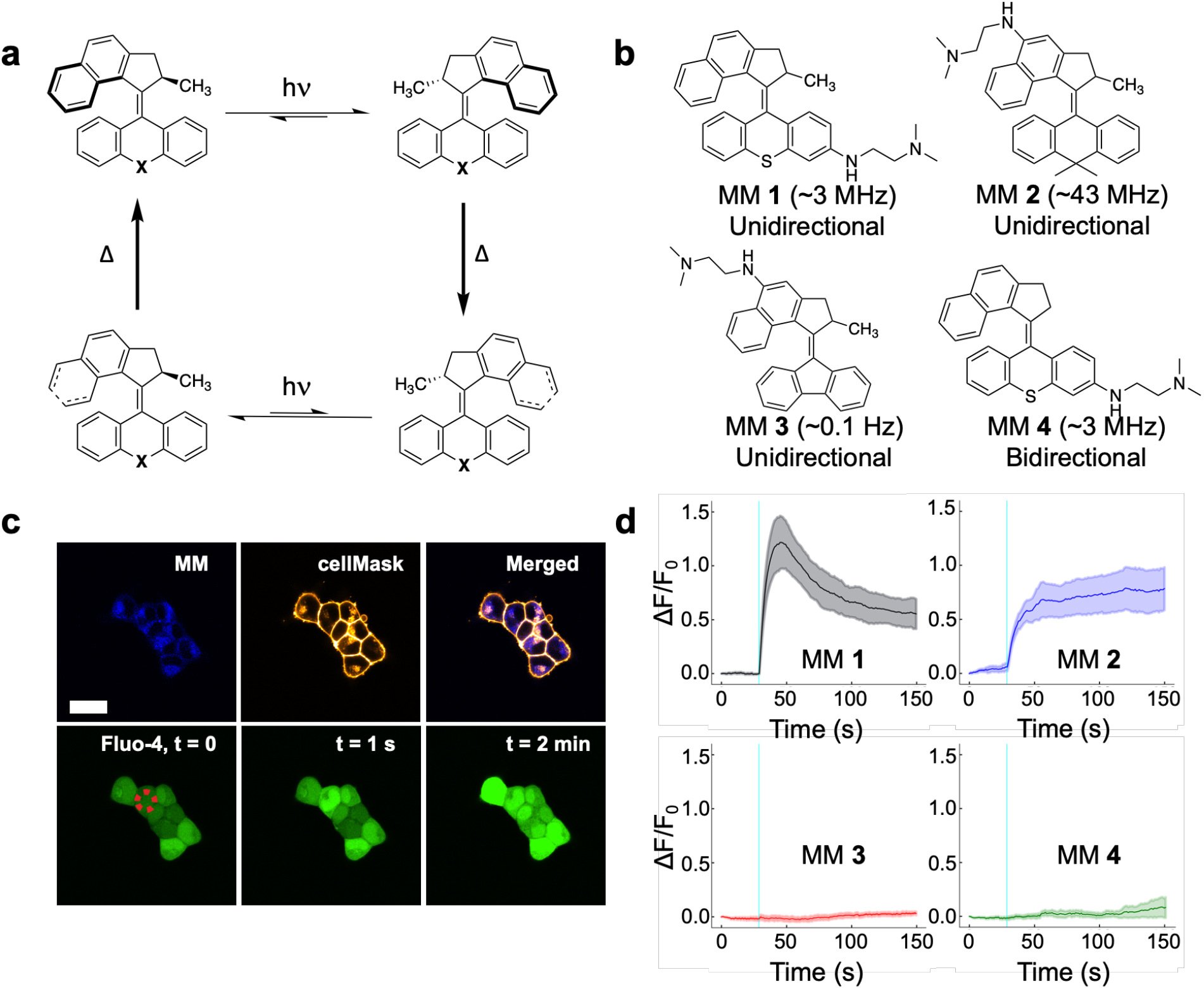
Structures of MM used in this study, their mechanism of rotation, and their activation to induce ICW in HEK293 cells. **(a)** The rotation cycle of a typical MM, showing two photoisomerizations (hν) and two thermal (Δ) helix inversions to complete a unidirectional rotation. The moiety in the “X” position can be changed to synthesize MM of different rotation speeds. **(b)** Structures of the MM employed in this study. MM **1** and MM **2** complete fast (MHz-scale) unidirectional rotation. MM **3** rotates at 0.1 Hz at room temperature. MM **4** lacks chirality imparted by the allylic methyl and possesses no preference for unidirectional rotation. **(c)** Confocal microscope images of cells treated with MM **1**, calcium-tracking dye Fluo-4, and cell membrane labeling dye CellMask (used to differentiate individual cells), showing a rapid increase in intracellular calcium levels after stimulation with 400 nm light. The red dotted circle labels the area of laser stimulation. The images in the top row were taken before stimulation, and the images in the bottom row were taken immediately prior to, 1 s after, and 2 min after stimulation. The scale bar applies to all images and is 20 μm. **(d)** Representative normalized fluorescence intensity traces of Fluo-4 in HEK293 cells treated with each MM. The solid line represents the average responses of n=6 independent cells, and the shaded area represents the standard error of the mean. MM **1**, MM **2**, and MM **4** were administered to cells at 8 μM. MM **3** was administered to cells at 24 μM. Stimuli for cells treated with MM **1, 2**, and **4** used a 250 ms pulse width delivered to a circular area of diameter 5 μm at 3.2×10^2^ W cm^-2^. Stimuli for cells treated with MM **3** were administered at 6.4×10^2^ W cm^-2^. For all plots, the cyan line indicates the time of stimulus presentation.

To investigate cellular responses to the actuation of MM, we employed live-cell calcium tracking. HEK293 cells were treated with the fluorescent intracellular calcium indicator Fluo-4 and loaded with MM. The structures of the MM employed in this study are shown in Fig. 1b, and the calcium responses of cells treated with fast-rotating MM **1** are shown in Fig. 1c. Stimulation of a single MM **1**-treated cell with a 400 nm laser (3.2×10^2^ W cm^-2^) increased Fluo-4 fluorescence in the targeted cell, reflecting a spike in the intracellular calcium concentration similar to that observed when cells are mechanically perturbed with a micropipette^8,18^. In the presence of the vehicle only, no calcium responses were evoked by the same laser treatment (Supplementary Fig. 1). Calcium responses were observed to propagate to adjacent cells (Supplementary Fig. 2) according to the degree of electrical connectivity between individual colonies. Similar responses were observed when cells were treated with scavengers of reactive oxygen species (ROS, Supplementary Fig. 3) and in experiments using X-Rhod-1 in place of Fluo-4 (Supplementary Fig. 4), suggesting that the observed responses do not depend on the production of ROS or fluorescence resonant energy transfer between MM **1** and calcium tracking dyes.

Cellular responses to MM were repeatable (Supplementary Fig. 5), and their amplitude could be controlled by the intensity of incident light (Supplementary Fig. 6). The strength of the evoked response also determines the downstream effects of stimulation. At typical stimulation irradiances (3.2×10^2^ W cm^-2^ for 250 ms), cells recovered from stimulation and showed no signs of apoptosis or necrosis (see Supplementary Fig. 7-8 and the accompanying discussion). Cells stimulated at higher intensities (6.4×10^2^ W cm^-2^ for 4 s) showed membrane blebbing and calcium accumulation over a 30 min period, indicating cell death (Supplementary Fig. 9)^25^. Hence, MM-induced ICW can be tuned between physiological, supraphysiological, and pathophysiological response regimes by adjusting the stimulus intensity. Calcium responses induced by MM **1** actuation were also observed in other cell lines, including neuroblast (N2A; Supplementary Fig. 10) and HeLa cells (Supplementary Fig. 11).

We investigated the calcium responses elicited by a library of MM consisting of two fast-rotating motors (MM **1-2**) and two complementary motors (MM **3-4**) used to test the effects of rotation speed and directionality (Fig. 1b). MM **1**, mimicking designs previously shown to kill cancer cells and antibiotic-resistant bacteria^26-28^, was synthesized with a six-membered ring stator containing a central sulfur atom. MM **1** rotates unidirectionally at ∼3 MHz. MM **2** and MM **3** are chemically similar to MM **1** but rotate at different rates. MM **2**, which rotates at ∼43 MHz, was synthesized with a six-member ring stator with two methyl groups branching off the center carbon atom^24^. MM **3**, a slow-rotating motor that rotates at ∼0.1 Hz^24^, was synthesized with a fluorene stator. Finally, MM **4** is an analog to MM **1** that lacks a stereogenic center at the allylic methyl site, which confers preference for unidirectional rotation. Without this stereogenic center, MM **4** “flaps” bidirectionally and switches stochastically between photoisomerization states. All four of these motors possess an appended aniline that shifts their absorption into the visible spectrum, enabling their observation by visible-light microscopy and the activation of their rotation with visible light^28^. These terminal amines are protonated at physiological pH, promoting their interaction with the hydrophilic heads of lipid bilayers in cell or organelle membranes.

Fig. 1d shows the calcium responses of cells treated with each MM and light. MM **3** was used at 3x concentration because of its lower extinction coefficient relative to the other MM (Supplementary Fig. 12, Supplementary Table 1). Fast-rotating MM **1** and MM **2** elicited rapid increases in intracellular calcium. MM **1** elicited high-amplitude transients that peak ∼10-20 s after stimulation and then decay over the next minute, whereas MM **2** elicited a more stable increase in intracellular calcium that does not decay as quickly as MM **1**. The different calcium release kinetics induced by MM **1** and MM **2** may be related to differences in their photoisomerization efficiency (see the discussion in the Supplemental Information)^23^. Meanwhile, slow-rotating MM **3** elicited no change in calcium activity upon irradiation, even at 3x the concentration and twice the stimulus intensity used to activate MM **1** and MM **2** (Fig. 1d). MM **4**, the fast-rotating motor with no preference for unidirectional rotation, elicited only small changes in calcium concentration. This experiment provides evidence to link MM rotation speed and directionality to their evoked cell signaling behavior.

### Mechanistic Study of MM-Induced ICW

Next, we studied the biological mechanism behind MM-driven calcium signaling. Calcium equilibrium in the cytosol is regulated by both export across the plasma membrane and uptake into the ER *via* membrane ATPases (Fig. 2a)^1^. Consequentially, the cytosolic concentration of calcium is typically low (∼100 nM) compared to those found inside the ER or extracellular medium (∼1.5 mM). Cytosolic calcium spikes commonly involve the entry of calcium from one of these two locations. Fluorescence microscopy showed that MM internalize within cells and interact with subcellular organelles, including mitochondria and the ER (Fig. 2b; Supplementary Figs. 13-14). Since MM are distributed primarily within cells, we hypothesized that MM-induced calcium responses arise from the release of calcium from intracellular stores. To test this hypothesis, we depleted calcium stores inside and outside of HEK293 cells and blocked various plasma membrane or ER calcium channels prior to stimulation (see Supplementary Table 2 for a complete list of the manipulations employed in these experiments).

**Fig. 2.**
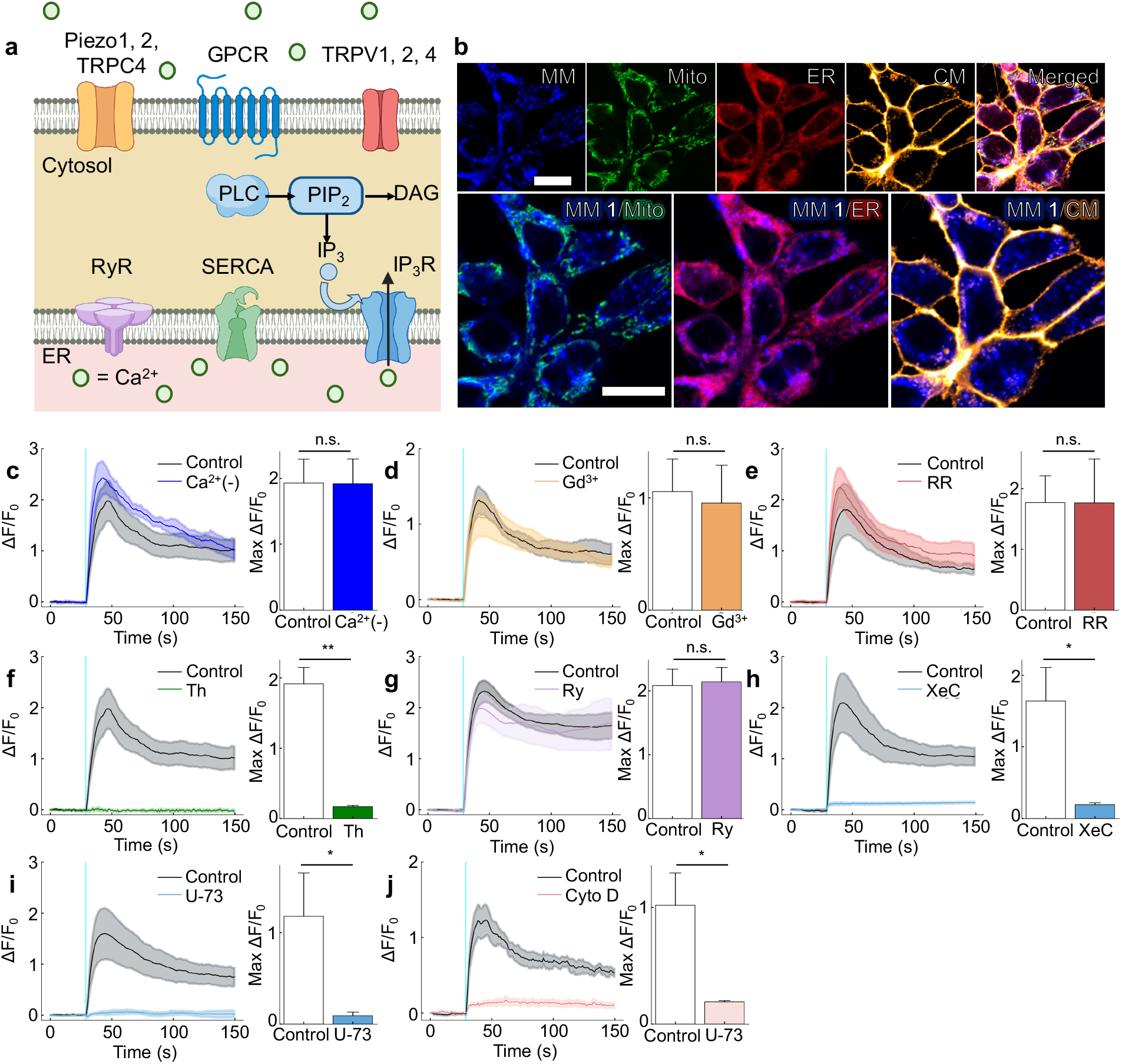
Mechanistic study of MM-induced ICW. **(a)** Schematic of possible mechanisms by which calcium can enter the cytoplasm. Prepared with Biorender.com. **(b)** Confocal microscope images of HEK293 cells treated with MM **1**, MitoTracker Green, ER Tracker Red, and CellMask Deep Red. Scale bars are both 20 μm and apply to the images in the same row. **(c-h)** Calcium waves elicited by MM **1**-treated cells **(c)** in calcium-free PBS, **(d)** in RR (1 μM), **(e)** in Gd^3+^ (50 μM), **(f)** after treatment of cells with Th (1 μM), **(g)** in Ry (100 μM), **(h)** in XeC (25 μM), **(i)** in U-73 (10 μM), **(j)** after pre-treatment of cells with Cyto D (2 μM). The solid line represents the calcium profiles averaged from six independent cells. The shaded region represents the standard error of the mean (n=6). Stimuli were presented after 30 s of imaging in each case, and the time of stimulus presentation is indicated by the vertical cyan line. For all plots, the black trace shows a positive control consisting of MM **1**-treated cells in typical imaging buffer recorded on the same day. All stimuli were delivered with a pulse width of 250 ms to a circular area of diameter 5 μm at a power ranging from 3.2×10^2^ W cm^-2^ to 5.1×10^2^ W cm^-2^. For all plots, the cyan line indicates the time of stimulus presentation. Controls and experimental groups were imaged and stimulated on the same day using the same conditions and with the same batch of cells. Error bars in the bar graphs represent the standard deviation of the mean of n=3 experiments (at least 6 stimulated cells per experiment). Statistical analyses were performed using a one-tailed Welch’s t-test. * *P*-value < 0.05, ** *P*-value < 0.01, *** *P*-value < 0.001, **** *P*-value < 0.0001.

Cells treated with MM **1** and stimulated by light pulses in the absence of extracellular calcium (Fig. 2c) showed no differences in response amplitude compared to a positive control in calcium-containing medium. Similar results were observed when plasma membrane calcium channels were blocked. Treatment of cells with ruthenium red (RR), a pharmacological inhibitor of temperature-sensitive vanilloid transient receptor potential (TRP) channels^29^, did not decrease the magnitude of MM-induced calcium responses (Fig. 2d). Similarly, treatment of cells with Gd^3+^, which is commonly used to block the effects of mechanosensitive plasma membrane channels such as Piezo1, Piezo2, and TRPC4^30^, also did not affect the observed responses (Fig. 2e). In all, our experiments show that plasma membrane TRP channels and extracellular calcium do not appreciably contribute to MM-induced calcium responses.

On the other hand, cells treated with MM **1** and thapsigargin, a sarco-endoplasmic reticulum calcium pump (SERCA) antagonist that depletes intracellular calcium^31^, did not show any measurable calcium flux upon light stimulation (Fig. 2f; *P* = 0.00257 < 0.01). This result implies that MM-evoked calcium responses arise from the release of ER-bound calcium stores by MM **1**.

Further mechanistic studies were conducted to determine how MM activation releases calcium from the ER. The mammalian ER predominantly expresses two large tetrameric calcium channels: IP_3_ receptors (IP_3_Rs) and ryanodine receptors (RyRs)^1,32^. In IP_3_-mediated calcium signaling, G-protein coupled receptors (commonly Gq/11 subtypes) expressed in the plasma membrane activate phospholipase Cβ (PLCβ) and tyrosine kinase receptors activate PLCγ to cleave phosphatidylinositol 4,5-biphosphate into diacylglycerol and IP_3_^32^. IP_3_ can then diffuse to the ER, where it binds to IP_3_R and causes calcium release. The activation of this network drives such a wide array of cellular activities that it is often referred to as simply “calcium release” due to its ubiquity in receptor biology^1^. RyRs are also expressed in many cell types and amplify existing calcium signals through calcium-induced calcium release.

Treatment of cells with ryanodine (Ry; 100 μM) to block RyR signaling has no effect on response amplitude (Fig. 2g) in HEK293 cells. However, treatment of cells with xestospongin C (XeC; 25 μM), a known antagonist of IP_3_R^33^, significantly diminished the strength of cellular responses to MM (Fig. 2h; *P* = 0.0167 < 0.05). Similar effects were observed when cells were treated with the PLC antagonist U-73122 (U-73; 10 μM) as an alternate method of blocking IP_3_ signaling (Fig. 2i; *P* = 0.0281 < 0.05). Furthermore, MM-induced responses were also inhibited in cells treated with cytochalasin D (Cyto D; 2 μM), an inhibitor of F-actin polymerization that disrupts PLC signaling by increasing the spatial distance between PLC and IP_3_R (Fig. 2j; *P* = 0.01762 < 0.05)^34^. These results implicate IP_3_ signaling as a primary driver of MM-induced calcium waves. The IP_3_ pathway is known to contribute to mechanosensitive calcium currents^35^ and is also involved in the induction of ICW in response to mechanical stimulation with a micropipette^2,18^.

### MM Cause Muscle Contraction In Cardiomyocytes

Next, we investigated whether MM-elicited ICW could be used to modulate calcium-driven biological processes, such as the beating activity and contraction of cultured cardiomyocytes. Fig. 3 shows the effects of MM stimulation on primary rat cardiomyocytes. MM distribute to the sarcoplasmic reticulum (SR; Fig. 3a), where subsequent light activation triggers localized calcium release (Fig. 3b, Supplementary Movie 1). MM-induced calcium release initially leads to localized myocyte contraction at the site of stimulation (Fig. 3c, kymograph 1, top arrow) likely due to the calcium-mediated activation of troponin and subsequent actin-myosin cross-bridge formation^36^. Then, SR calcium release induces beating in quiescent cardiomyocytes and accelerated beating in active cardiomyocytes (Fig. 3c, bottom arrow in kymograph 1 and arrow in kymograph 2; Supplementary Movie 2).

**Fig. 3.**
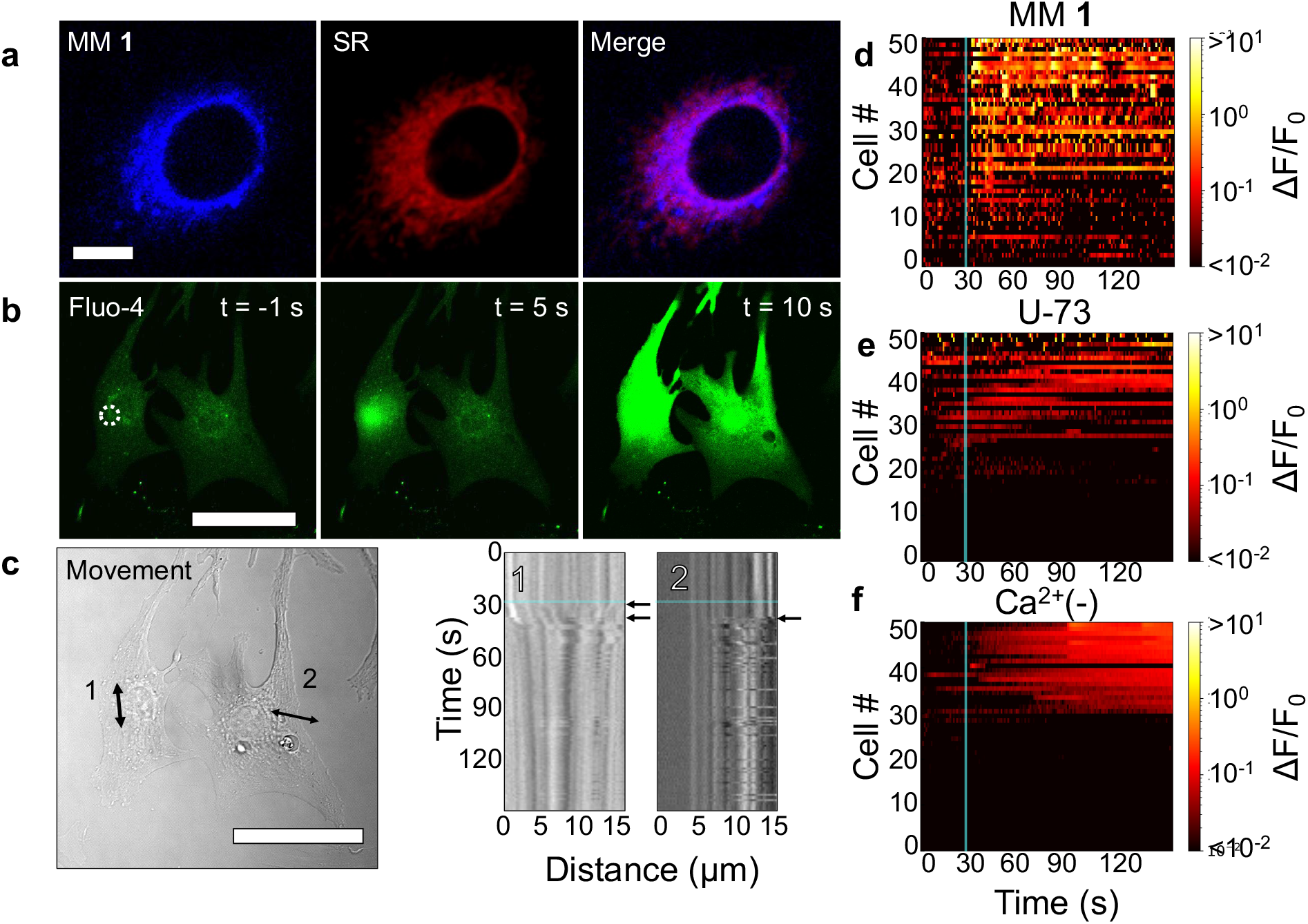
MM-induced calcium waves cause localized calcium release, contraction, and beating in cardiomyocytes. **(a)** Confocal microscope images of a single myocyte treated with MM **1** and ER Tracker Red. The scale bar is 10 μm and applies to all images. (**b)** Confocal microscope images showing Fluo-4 fluorescence, revealing calcium activity in cardiomyocytes before and after stimulation. The scale bar is 50 μm and applies to all images. The white circle represents the stimulation region. **(c)** Kymographs representative of cardiac myocyte contractile responses from the same cells shown in (b). The line profiles from which kymographs **1** and **2** were taken are shown as double-sided arrows in the bright field image. Kymographs **1** and **2** were taken from the stimulated cell and an adjacent cell, respectively. The scale bar in the bright field image is 50 μm. Kymograph **1** shows local contraction in the stimulated cell (top arrow in **1**), indicated by a convergence of the edges in the kymograph plot. Both kymographs **1** and **2** show periodic beating in the stimulated cell and surrounding cell (bottom arrow in **1** and arrow in **2**). **(d-f)** Normalized fluorescence intensity change of Fluo-4 in cardiomyocytes adjacent to stimulated cardiomyocytes that were treated with **(d)** MM **1** alone, **(e)** MM **1** and U-73 (10 μM), and **(f)** MM **1** in calcium-free PBS. For each condition, n=50 cells were used. The x-axis label in (f) also applies to (d-e). All stimulation experiments were performed with a stimulation time of 250 ms at 5.1×10^2^ W cm^-2^ in a circular region of 5 μm diameter using a 400 nm laser. In all plots, the cyan line indicates the time of stimulus presentation.

We tracked the behavior of colonies of cardiomyocytes in contact with cells stimulated with MM and light to determine whether we could use MM to drive biological behaviors coordinated in networks of cells, such as contraction. Colonies of cardiomyocytes adjacent to stimulated cells responded to stimulation by firing action potentials (APs) or participating in the generated calcium wave (Fig. 3d; Supplementary Figs. 15-16). Activation of adjacent cardiomyocytes in response to stimulation could be prevented by either inhibiting IP_3_-mediated calcium release in the stimulated cell (Fig. 3e) or by preventing the influx of calcium from outside the cell during AP firing (Fig. 3f). Cells stimulated with MM **1** and light in calcium-containing medium exhibited firing rates of 5.1 spikes min^-1^ cell^-1^, whereas cells stimulated with MM **1** and light in the presence of PLC inhibitor (U-73; 10 μM) exhibited firing rates of ∼0.8 spikes min^-1^ cell^-1^ (*P* < 0.0001 by a one-tailed Welch’s t-test). Meanwhile, cells stimulated in the absence of extracellular calcium did not exhibit any spiking activity, suggesting that the increased activity of cardiomyocyte colonies in response to MM may be due to the action of voltage-gated ion channels in the plasma membrane of myocytes. These ion channels are likely triggered by local depolarization of the membrane induced by SR calcium release^37^. These experiments show that biological behaviors coordinated in networks of cells, such as contraction, can be controlled by intracellular MM-induced calcium wave generation.

### MM Control Behavior In Vivo

Finally, using an *in vivo* model of muscle contraction, we sought to investigate whether MM-induced ICW can control biological activity at the organism level. For this purpose, we chose *Hydra vulgaris* as a model system. *Hydra* are radially symmetric, millimeter-sized freshwater cnidarians containing tentacles, an oral region, and an aboral region connected by a long, tubular body column. In the oral region, *Hydra* have a dome-shaped structure called a “hypostome” surrounded by a ring of tentacles. At the other extremity, they have a foot called the “peduncle,” which the animal uses to attach to substrates (Fig. 4a). *Hydra* were chosen as a model system because of their small size, lack of chitin layer, excitable epitheliomuscular tissue, and simple nervous system. In addition, *Hydra* exhibit spontaneous and stimulus-controlled contractions, both driven by ICW^38^, which can be directly visualized due to their transparent body. In our experiments, we used *Hydra* lines genetically engineered to express the calcium indicator GCaMP7b in their endothelial epitheliomuscular tissue (see *Methods*).

**Fig. 4.**
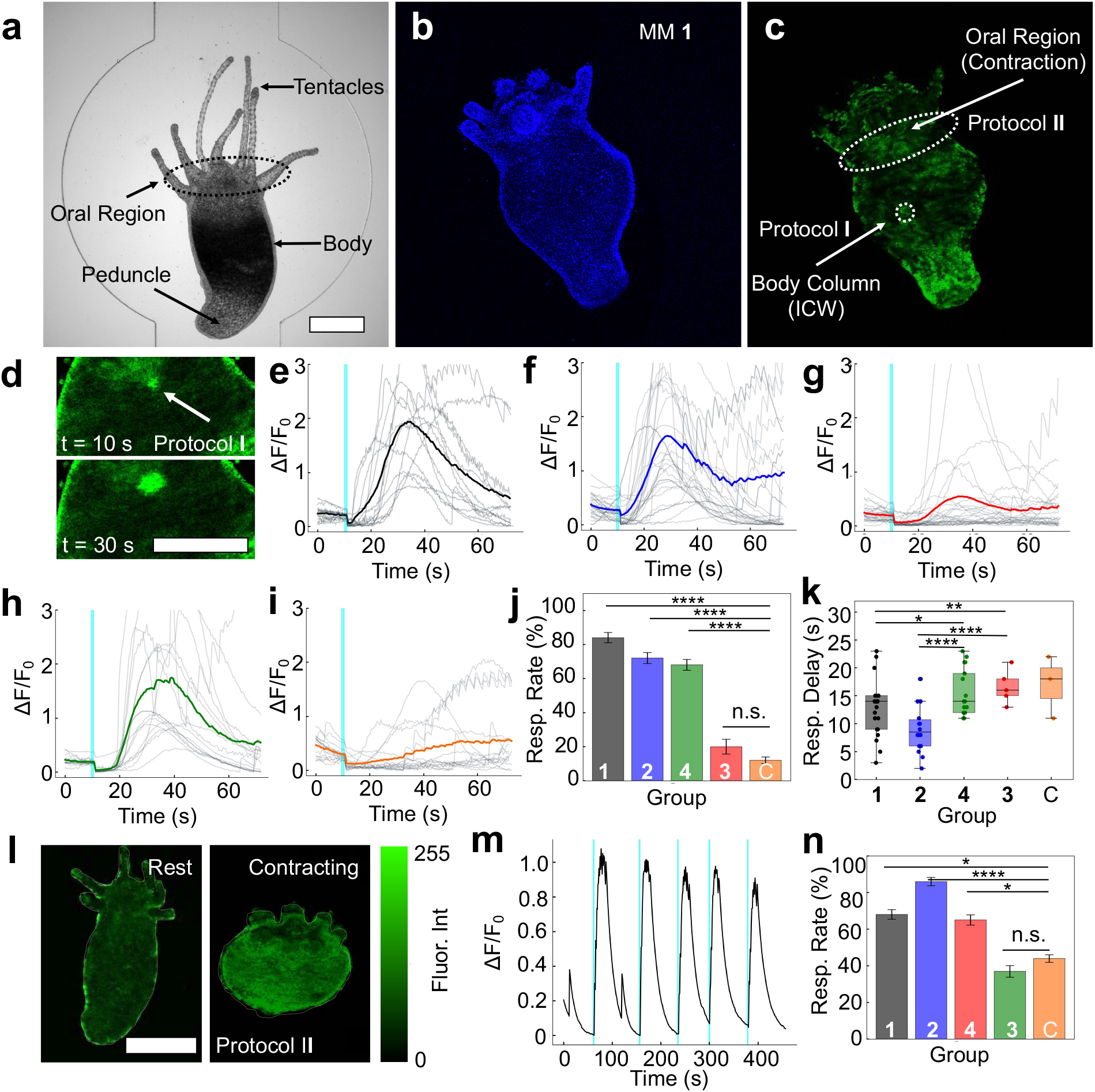
MM induce ICW and muscle contraction *in vivo*. **(a)** Image of a *Hydra* loaded into a microfluidic chamber with anatomical regions marked. Scale bar is 100 μm and applies to (a-c). **(b)** MM **1** (24 μM) loaded into *Hydra* for 24 h. **(c)** A *Hydra* expressing GCaMP7b in endodermal epitheliomuscular cells. The stimulation regions used for Protocols **I** and **II** are marked. Protocol **I** was used for body column stimulation (1 s pulse to a 10 μm diameter circular area) to trigger regional ICW. Protocol **II** was used for oral region stimulation (2 s pulse to the oral region, ∼1000-2000 μm^2^) to trigger whole-body contraction. **(d)** A regional ICW in a *Hydra* treated with MM **1** and stimulated with 405-nm light via Protocol **I**. The indicated times represent the time after stimulation at which the image was taken. Scale bar applies to both images and is 100 μm. **(e-i)** Responses observed from *Hydra* treated with **(e)** MM **1, (f)** MM **2, (g)** MM **3, (h)** MM **4**, or **(i)** DMSO control (“C”) and stimulated for 1 s using Protocol **I**. Bold, colored traces represent the average trace (n=25 across 5 *Hydra*). Gray traces represent individual experiments. **(j)** Bar graph showing ICW response rate and **(k)** box-and-whisker plot showing ICW delay time of *Hydra* stimulated using Protocol **I** across MM treatment conditions (n=25 experiments across at least 5 *Hydra* per condition). ICW responses were defined as ΔF/*F*_0_ > 1. **(l)** Representative contraction in a *Hydra* treated with MM **2** and stimulated via Protocol **II**. Scale bar applies to both images and is 100 μm. **(m)** Representative GCaMP7b fluorescence trace in a *Hydra* treated with MM **2** and stimulated using Protocol **II. (n)** Bar graph showing contractile response rates across treatment conditions for *Hydra* stimulated for 2 s using Protocol **II** (at least 50 experiments across ≥5 *Hydra* per condition). Error bars in bar graphs represent the standard error of the mean. See *Methods* for more info on the calculation of ICW and contractile response rates. For all plots, cyan lines indicate the time of stimulus with 405-nm light (9.0×10^2^ W cm^-2^), during which no data was collected. * *P*-value < 0.05, ** *P*-value < 0.01, *** *P*-value < 0.001, **** *P*-value < 0.0001.

Prior to stimulation experiments, *Hydra* were loaded with MM by incubating with solutions containing 24 μM of MM for 24 h (Fig. 4b). Distinct stimulation protocols were employed to cause either local ICW or whole-body contraction (Fig. 4c). First, treatment of MM-loaded *Hydra* with pulses of laser light administered to a small region of the body column (Protocol **I**) caused ICW emanating from the site of stimulation (Fig. 4d; Supplementary Fig. 17; Supplementary Movies 3-4). Similar to ICW evoked *in vitro*, these ICW propagated throughout the *Hydra* body column according to the degree of electrical connectivity of the stimulated cells. Distinct propagation kinetics were observed when stimulating different regions of *Hydra* simultaneously (Supplementary Movie 5). Second, we attempted to use MM activation to drive whole-body *Hydra* contractions by administering laser stimuli to the oral region (Protocol **II**). We decided to target the oral region because mechanical stimulation of this region has been shown to stimulate burst contraction, likely *via* the sensory neurons that cluster in this region^39-41^. When laser stimuli were delivered to the oral region of *Hydra* treated with fast-rotating MM **1** and **2** (Protocol **II**), they exhibited contraction bursts associated with whole-body calcium waves (Supplementary Movie 6) similar to those observed with macro-mechanical stimulation^42^.

Fast-rotating MM were generally more successful in eliciting both regional ICW (Protocol **I**) and whole-body contractions (Protocol **II**). Fig. 4e-i show responses from MM-treated *Hydra* when stimulated *via* Protocol **I** to elicit regional ICW. Furthermore, Fig. 4j compares the response rates of *Hydra* treated with each MM for exhibiting regional ICW. Fast-rotating MM, including MM **1**, MM **2**, and even non-unidirectional fast motor MM **4**, consistently elicited robust regional ICW upon stimulation of the body column (Protocol **I**). Occasional responses were also observed from *Hydra* treated with slow-rotating MM **3**, but the response rate of these *Hydra* was not significantly different from solvent-only controls (Fig. 4j). *Hydra* also demonstrated marked differences in response kinetics depending on the type of MM employed. Fast, unidirectionally rotating MM elicited appreciably quicker responses (12.76 s for MM **1** and 9.05 s for MM **2**) than slow or non-unidirectionally rotating MM (15.46 s for MM **3** and 16.6 s for MM **4**; Fig. 4k).

Similar trends were observed for whole-body contractile response rates across *Hydra* treated with different MM and light conditions (see *Methods* for a detailed description of the response rate calculation). Fig. 4l-m show a representative whole-body contraction and GCaMP7b fluorescence trace from a typical experiment using MM **2** and stimulation using Protocol **II**. In these experiments, the fastest-rotating MM, MM **2**, was most successful at inducing *Hydra* contraction, demonstrating a response rate of 86% (Fig. 4n). This fast-rotating MM elicited contraction of *Hydra* at a higher rate than light alone (*P* < 0.00001) or slower-rotating MM **3** (*P* < 0.00001) by Fisher’s Exact Test. MM **1** also elicited contraction at a significantly higher rate than light alone (*P* = 0.0199 < 0.05), while the slow-rotating MM **3** (*P* = 0.5325 > 0.05) did not elicit contraction at a significantly higher rate than light alone. Interestingly, MM **4**, which rotates quickly but non-unidirectionally, elicited *Hydra* contraction with a response rate of 65% and was significantly (*P* = 0.0439 < 0.05) more effective than light alone. MM **4** also elicited regional ICW in *Hydra* despite causing only weak responses *in vitro*. These results imply that even the smaller responses elicited by MM **4** *in vitro* can be amplified *in vivo* across networks of cells. Consequentially, rotor speed is a better indicator of the ability of MM to drive *Hydra* contraction than rotor unidirectionality. Further experiments are needed to elucidate both the biological machinery responsible for amplifying MM-induced signals and the factors influencing MM propensity for causing ICW in *Hydra* (see the discussion in the Supplemental Information).

The contractile behavior of *Hydra* in relation to the presentation of stimulus is detailed in Supplementary Figs. 18-22. *Hydra* appeared to exhibit a photic response to light in the absence of MM (Supplementary Fig. 22; *P* = 0.0525 > 0.05) consistent with the photosensitivity of the hypostome and tentacles described previously^43^. However, fast-rotating MM actuation was notably superior at driving muscle contraction compared to light alone. Peak identification algorithms were employed to track calcium spikes in the fluorescence data and contraction bursts across the timescale of our experiments. In *Hydra* treated with MM **2** and light, contraction onset occurs predominantly upon the presentation of the stimulus, and a high density of contraction bursts appears 5-10 seconds later (Supplementary Fig. 19). In the absence of MM, this relationship weakens dramatically. In the absence of light, it disappears completely (Supplementary Fig. 22). These results suggest that the actuation of fast-rotating MM can control the behaviors exhibited by *Hydra* over the time scale of our experiments.

## Conclusion

This work demonstrated that the activation of MM triggers calcium waves in a fashion that depends on their fast, unidirectional rotation. MM localized to the endoplasmic reticulum, where their activation with sub-second pulses of visible light caused a rapid rise in intracellular calcium that propagated to neighboring cells. These ICW were shown to arise from IP_3_ signaling. The amplitude of MM-induced ICW was tunable by the laser light intensity. MM-induced ICW remotely triggered muscle contraction *in vitro* and *in vivo*, suggesting that MM can be used for molecular-scale mechanical control of biological activity.

## Experimental Methods

### Synthetic Chemistry

Details on the synthesis and characterization of MM **2-4** are provided in the Supplementary Information (Materials & Methods; Supplementary Figs. 23-28). Synthesis and characterization information on MM **1** is provided elsewhere^28^. MM were dissolved in DMSO at a concentration of 8 mM and sonicated for 5 s prior to use. MM solutions were stored at −20 °C in aluminum foil-wrapped containers to avoid degradation. UV-Vis spectra of MM were taken in spectral-grade water using a Shimadzu UV-3600 Plus spectrophotometer. Extinction coefficients were calculated by constructing a Beer’s Law plot using concentrations between 8 and 32 μM.

### Cell Culture and Preparation of Cells for Microscopy

HEK293 cells were chosen as the principal model system due to their widespread use in electrophysiological studies. HEK cells, like most excitable or non-excitable cells, are known to exhibit calcium responses^44^. HEK293 cells, HeLa cells, and N2A cells were cultured in DMEM (Lonza) supplemented with 10% FBS (Gibco), and 1% penicillin/streptomycin (Lonza) at 37 °C in 5% CO_2_ atmospheric conditions. Cells were passaged at <90% confluence. To grow cardiomyocytes, a dissolved rat heart (TransnetYX^©^) was triturated, centrifuged, and resuspended in cardiomyocyte growth medium (TransnetYX^©^ catalog #SKU-NBCG) and seeded at a concentration of 60,000 cells cm^-2^. Growth substrates were pre-treated with 1% gelatin solution for 3 h, then washed with PBS. Cardiomyocytes were allowed to grow for 4 days, after which the growth medium was exchanged for cardiomyocyte maintenance medium (TransnetYX^©^ catalog #SKU-NBCM). Experiments were conducted from day 4 to day 6.

Imaging was performed in imaging extracellular buffer (iECB; 119 mM NaCl, 5 mM KCl, 10 mM HEPES, 2 mM CaCl_2_, 1 mM MgCl_2_ (pH 7.2); 320 mOsm). Cells were prepared for imaging by seeding a 35 mm Ibidi imaging dish with ∼50,000 cells in 1 mL of complete growth medium, and grown for 2 days. Prior to imaging, cells were incubated with dyes and/or molecules resuspended in complete growth medium at the appropriate concentration [MM **1** (8 μM), MM **2** (8 μM), MM **3** (24 μM), MM **4** (8 μM), Fluo-4 (2 μM)]. MM and Fluo-4 were incubated with cells for 45 min. Unless otherwise specified, all experiments were performed in triplicate.

### In vitro Imaging and Stimulation

Cells were imaged with a Nikon A1-Rsi confocal system mounted on a widefield Ti-E fluorescence microscope. Imaging was performed using a 60x water immersion objective (NA of 1.27, 0.17 mm working distance). Green fluorophores (Fluo-4, MitoTracker Green) were excited with a 488 nm photodiode laser. Red fluorophores (ER-Tracker-Red, PI) were excited with a 561 nm photodiode laser. Deep red fluorophores (CellMask plasma membrane stain) were excited with a 630 nm photodiode laser. Laser stimulation for *in vitro* experiments was performed with a 400 nm photodiode laser (Coherent OBIS™ LX SF) operating in a fluorescence-recovery-after-photobleaching (FRAP) experiment mode, delivering up to 6.4×10^2^ W cm^-2^ at sample level. *In vitro* experiments with HEK293 cells were conducted using a stimulus irradiance of 3.2×10^2^ W cm^-2^ except where otherwise indicated. Cardiomyocyte experiments were conducted using a stimulus irradiance of 5.1×10^2^ W cm^-2^. Power was calibrated using a Thor Labs S130C laser power meter. Stimulation was targeted to a circular area of diameter 5 μm in a 250 ms pulse, during which the laser rastered across the entire region of interest. An additional laser stimulation set-up was used to verify the requisite stimulation power and demonstrate excitation in a non-scanning laser mode (see Supplementary Fig. 29 and the accompanying discussion). Fluo-4 fluorescence was collected for ∼30 s before and 2 min after stimulation (since the recorded time of the first data point is 0 s, stimulation occurs between 28.6 and 28.85 s in recorded traces). Images were collected using a Galvano scanner operating at 0.94 fps. In some experiments, images were also collected using a resonant scanner operating at 7.7 fps.

### Colocalization Analysis

Cells were treated with MitoTracker Green (ThermoFisher, 400 nM), ER Tracker Red (ThermoFisher, 500 nM), and CellMask Deep Red Plasma Membrane Stain (ThermoFisher, 500 nM). ER Tracker Red and MitoTracker Green were loaded into cells for 45 min in complete growth medium. Then, CellMask Deep Red Plasma Membrane Stain was loaded into cells for 5 min and cells were subsequently loaded into the microscope chamber. Images for co-localization analysis were collected in a Nikon A1 confocal microscope using a 60x water immersion objective. Z-stack images spanning a minimum 20 μm range were collected and processed using the Coloc-2 plugin in Fiji. Colocalized pixel intensity maps were generated using the colocalization threshold function in Fiji.

### Pharmacological Experiments

In pharmacological blocking experiments, cells were loaded with MM and Fluo-4 as previously described. Typical imaging experiments were then performed on six cells in iECB alone as a positive control. Then, the imaging buffer was replaced, and the cells were treated as required for each experiment (see below). After a brief recovery period, six non-previously imaged cells were stimulated and imaged. Phosphate-buffered saline (PBS) was used for experiments in calcium-free buffer. Thapsigargin (Th; 1 μM) and cytochalasin D (Cyto D; 2 μM) were incubated with cells in the incubator in complete growth medium for 1 h (Th) or 2 h (Cyto D) before imaging. Ruthenium red (RR; 1 μM), gadolinium (Gd^3+^; 50 μM), ryanodine (Ry; 100 μM), xestospongin C (XeC; 20 μM), and U-73122 (U-73; 10 μM) were administered directly into the iECB after control imaging was finished. Cells were incubated in the medium for ∼5 min prior to imaging. For experiments using ROS scavengers, cells were incubated with either melatonin (Mel; 100 μM), thiourea (thioU; 50 mM), or L-ascorbic acid (VitC; 2 mM) for 1 h prior to experiments. The traces shown are representative results across at least six cells from individual experiments, while bar graphs represent the average results across at least three distinct experiments. See Supplementary Table 2 for more information.

### Hydra Preparation

*Hydra* were grown in *Hydra* medium containing CaCl_2_·2H_2_O (1 mM), MgCl_2_·6H_2_O (0.1 mM), KNO_3_ (0.03 mM), NaHCO_3_ (0.5 mM), and MgSO_4_ (0.08 mM) in deionized water at 18 °C in a light-cycled (12 h light, 12 h dark) incubator. *Hydra* were fed with an excess of freshly brined *Artemia Nauplii* (Brine Shrimp Direct, Ogden, UT, # BSEP 8Z) three times a week. All experiments were performed at room temperature after starving the animals for 2 days. The transgenic line expressing calcium indicator GCaMP7b under the Ef1α promoter in endodermal epitheliomuscular tissue was generated by embryonic microinjection in a collaboration by the Robinson lab (Rice University) and the Juliano lab (University of California, Davis)^42^.

### Hydra Imaging and Stimulation

*Hydra* were incubated with the selected MM for 24 h before imaging. Stimuli were applied using an ROI-driven fluorescence-recovery-after-photobleaching (FRAP) mode in a Nikon A1 confocal microscope. All *Hydra* were able to recover after stimulation experiments, indicating both MM administration and subsequent stimulation treatments were not toxic to the *Hydra*.

Distinct stimulation protocols were employed to elicit either regional ICW or whole-body contraction. In Protocol **I**, stimulation was delivered using a 405-nm laser diode at 9.0×10^2^ W cm^-2^ in 1 s pulses delivered to a 10 μm region of the *Hydra* body column. Protocol **I** resulted in regional ICW. In Protocol **II**, stimulation was delivered using a 405-nm laser diode at 9.0×10^2^ W cm^-2^ in a 2 s pulse delivered to the oral region of the *Hydra* (typical area of ∼1,000-2,000 μm^2^). Protocol **II** resulted in whole-body calcium waves and contraction. Note that despite the longer stimulation time used in Protocol **II**, the *Hydra* still experiences less pixel dwell time and less incident light per unit area compared to *in vitro* experiments and Protocol **I** due to the size of the stimulation region (∼50-100x larger depending on the size of the animal). GCaMP7b fluorescence were recorded using a 456 nm laser diode for fluorophore excitation. In experiments using Protocol **I**, stimuli were presented at 10 s. In experiments using Protocol **II**, stimuli were presented at irregular intervals at least 60 s apart to prevent interference from periodic spontaneous contractions of *Hydra*.

### Data Analysis

Data were analyzed using custom-written Python scripts. Fluorescence traces of the calcium indicator Fluo-4 from cells were imported and processed using the Pandas library. F^0^ was calculated as the average fluorescence intensity in the first 10 frames (∼10 s) of the acquisition and used to calculate ΔF/*F*_0_ over the entire length of the acquisition. Since data was not collected during stimulation, a “dead time” equivalent to the time of stimulation was manually added to each recording. Spiking behavior of cardiomyocytes was calculated using the SciPy peak finder function.

Fluorescence traces of calcium indicator GCaMP7b in *Hydra vulgaris* were processed similarly to *in vitro* data and baseline-corrected to set the minimum measured ΔF/*F*_0_ value as “0” due to the spontaneous, periodic activity of the *Hydra*. For experiments analyzing regional ICW (stimulating using Protocol **I**), responses reaching ΔF/*F*_0_ > 1 in the stimulated area were considered a “success” in the calculation of the ICW response rate. The “delay time” was the time at which ΔF/*F*_0_ reached 1. Peaks corresponding to whole-body contraction were not included in the analysis of ICW amplitude. ICW response rate was calculated across n=25 experiments for each condition across at least 5 *Hydra*.

In experiments analyzing whole-body contraction (stimulating using Protocol **II**), the induction of a contractile “response” comprising a change in body GCaMP7b fluorescence of ΔF/*F*_0_ > 1.5x within 3 s of stimulus presentation was counted as a “success” in the calculation of the contractile response rate. To identify contraction onset, a limit-of-detection peak finding algorithm was used with a threshold value of 1.5. To identify individual calcium spikes in a contraction burst, the SciPy peak finder function was employed. Induction of a contractile “response” comprising a change in body GCaMP7b fluorescence of ΔF/*F*_0_ > 1.5x within 3 s of stimulus presentation was considered a “success” when calculating response rates. Response rates were calculated over at least 50 presentations of stimuli. Stimuli presented at a time when the *Hydra* were already contracting (ΔF/*F*_0_ > 0.3) were discarded. *Hydra* that exhibited mouth-opening behavior during the experiment were discarded. The effects of spontaneous and photic responses were accounted for by comparison with DMSO and DMSO and light controls. For more information on *Hydra* data processing, see Supplementary Table 3.

### Statistical Analysis

One-tailed Welch’s t-tests were performed to assess differences in cellular responses after pharmacological manipulations, or when treated with MM of different rotation speeds. Fisher’s Exact Tests was used to assess differences in contractile response rates between *Hydra vulgaris* treated with MM of variable rotation speed and light. * *P*-value < 0.05, ** *P*-value < 0.01, *** *P*-value < 0.001, **** *P*-value < 0.0001.

## Acknowledgements

This project received funding from the Discovery Institute, the Robert A. Welch Foundation (C-2017-20190330), the National Science Foundation Graduate Research Fellowship Program (JLB), the DEVCOM Army Research Laboratory under Cooperative Agreement W911NF-18-2-0234 (AVV), and the European Union’s Horizon 2020 research and innovation programme under the Marie Sklodowska-Curie grant agreement No. 843116 (ALS). The views and conclusions contained in this document are those of the authors and should not be interpreted as representing the official policies, either expressed or implied, of the Army Research Laboratory or the U.S. Government. The U.S. Government is authorized to reproduce and distribute reprints for Government purposes notwithstanding any copyright notation herein. The funders had no role in the study design, data collection and analysis, decision to publish, or preparation of the manuscript. This work was conducted in part using resources from the Light Microscopy Facility and the Shared Equipment Authority at Rice University. The authors acknowledge Zachary C. Sanchez (Vanderbilt University) for useful help and advice regarding growth and activity of cardiomyocytes.

## Author Contributions

Conceptualization, JLB, ARV, JMT.

Methodology, JLB, GD, SK, JZ, ALS.

Organic synthesis: ARV, GL, BL, JMT.

Formal analysis, JLB.

Investigation, JLB, SK, GC, DA, JZ.

Resources, JTR, JMT.

Writing, Original draft, JLB.

Writing, Reviewing, and Editing, JLB, GD, SK, ALS, JTR, JMT.

Visualization, JLB.

Supervision, GD, JTR, JMT.

Funding acquisition, JMT.

Project oversight: JMT.

## Biological materials

All biological materials used in this work are available from commercial sources.

## Data availability

All data supporting the findings of this study are available within the article and its supplementary information files.

## Code availability

Data analysis scripts are available upon request to the author, JMT.

