## Supplemental Information for "Molecular Machines Stimulate Intercellular Calcium Waves and Cause Muscle Contraction"

A. L. Santos

Department of Chemistry, Rice University, 6100 Main Street MS 222, Houston, TX 77005 USA.

IdISBA - Fundación de Investigación Sanitaria de las Islas Baleares, Palma, Spain

Associate Professor J. T. Robinson

Department of Bioengineering, Department of Electrical Engineering, Rice University, 6500 Main Street BRC 973, Houston, TX 77005 USA.

Professor J. M. Tour

Department of Chemistry, Smalley-Curl Institute, NanoCarbon Center, Welch Institute for Advanced Materials, Department of Materials Science and Nanoengineering, Department of Computer Science, Rice University, 6100 Main Street MS 222, Houston, TX 77005 USA.

### Materials and Methods

Synthesis of MM **2**

Final product characterization of MM **2** is reported in Supplementary Figs. 23-24.

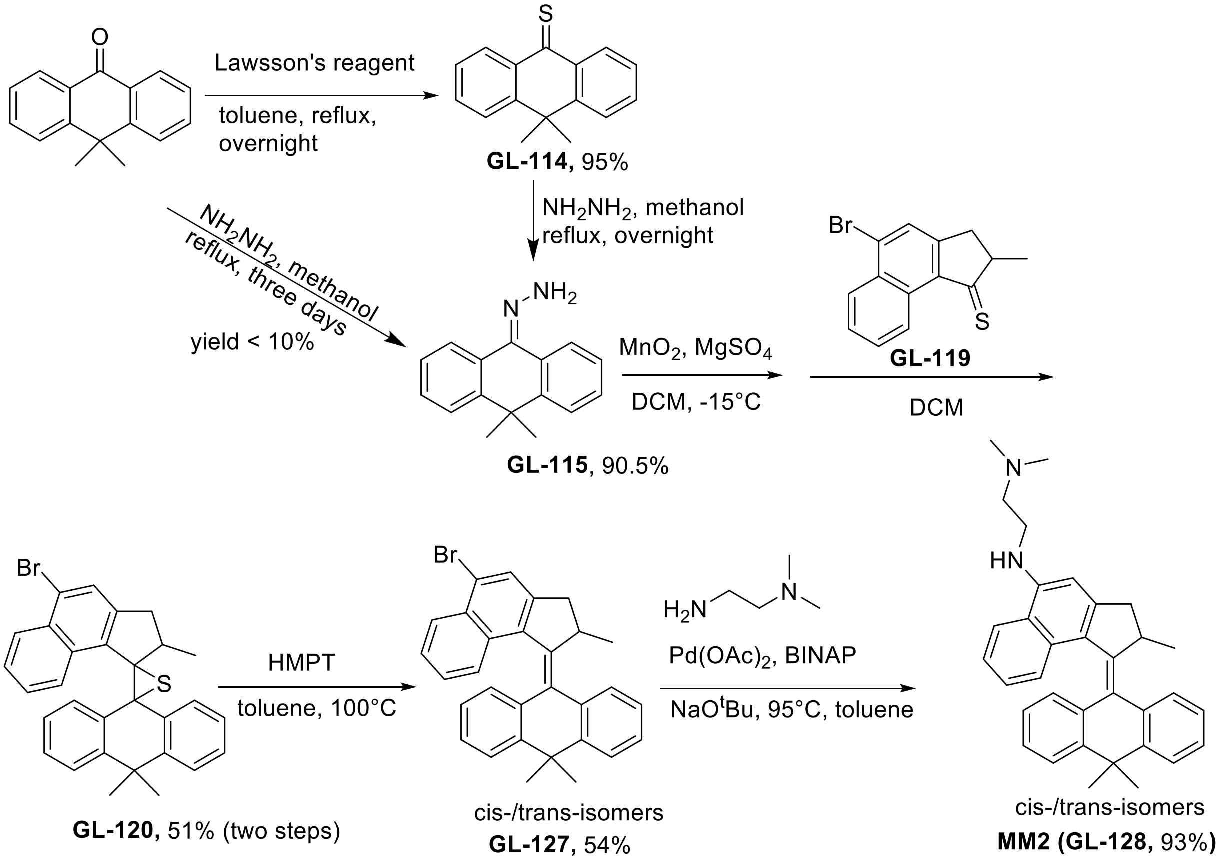


**Supplementary Scheme 1.** Synthesis of MM **2** (herein denoted GL-**128**).

**General Methods.** All glassware was oven-dried prior to use. Reagent grade dichloromethane (DCM, $CH_2Cl_2$) was distilled from calcium hydride ($CaH_2$) under a nitrogen atmosphere. All reactions were carried out under a nitrogen atmosphere unless otherwise noted. All other chemicals were purchased from commercial suppliers and used without further purification. Flash column chromatography was performed using 230-400 mesh silica gel from EM Science. $^{1}H$ NMR and $^{13}C$ NMR spectra were recorded at 600 and 150 MHz, respectively. Chemical shifts (δ) are reported in ppm from tetramethylsilane (TMS). Compound GL-**119** was synthesized according to the previous literature[28].

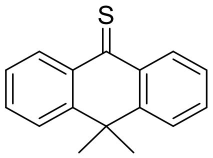

**GL-114**

10,10-Dimethyl-9(10H)-anthracenone (1.67 g, 7.5 mmol) and Lawesson's reagent (6.08 g, 15 mmol) were mixed in a solution of toluene (30 mL). The solution was heated to reflux for 14 h. The solution was permitted to cool to room temperature. The toluene was removed by rotary evaporation. The residue was purified by silica gel chromatography using ether/hexanes (1/4), affording the title compound GL-**114** as a blue solid in 95% yield (1.58 g). FTIR (KBr): 3060, 2974, 2928, 2862, 1723, 1594, 1578, 1558, 1478, 1443, 1310, 1301, 1277, 1226, 1204, 1165, 1023, 903, 785, 759, 637 $cm^{-1}$. $^{1}H$ NMR (600 MHz, $CDCl_3$) δ 8.61 (m, 2H, Ar-*H*), 7.64 (m, 4H, Ar-*H*), 7.35 (m, 2H, Ar-H), 1.72 (s, 6H, C*H₃*) ppm. $^{13}C$ NMR (150.8 MHz, $CDCl_3$) δ 144.4, 138.0, 132.9, 130.3, 126.7, 126.6, 126.1, 125.6, 38.9, 32.9, 32.1 ppm. HRMS (ESI) calcd for $C_{16}H_{15}S$ $[M+H]^+$: 239.0894. Found: 239.0901.

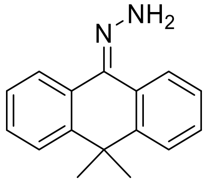

**GL-115**

Hydrazine (20 mL) was added into a solution of **GL-114** (1.36 g, 5.71 mmol) in methanol (30 mL) in a round-bottom flask. The solution was heated to reflux for 16 h. After permitting to cool to room temperature, the solvents were removed by rotary evaporation. The residue was purified by silica gel chromatography using ether/hexanes (1/4) as eluent, affording the title compound **GL-115** in 90.5% yield (1.28 g). FTIR (KBr): 3389, 3206, 3064, 3029, 2971, 2922, 1658, 1622, 1597, 1570, 1468, 1447, 1385, 1360, 1343, 1291, 1264, 1185, 1094, 1040, 944, 765, 734, 654 $cm^{-1}$. $^1H$ NMR (600 MHz, $CDCl_3$) δ 8.18 (m, 1H, Ar-*H*), 8.02 (m, 1H, Ar-*H*), 7.35 (m, 1H, Ar-*H*), 7.27 (m, 1H, Ar-*H*), 7.14-7.09 (m, 2H, Ar-*H*), 7.06-7.03 (m, 2H, Ar-*H*), 5.35 (s, 2H, N*H*$_2$), 1.40 (s, 6H, C*H*$_3$) ppm. $^{13}C$ NMR (150.8 MHz, $CDCl_3$) δ 148.2, 143.9, 141.1, 136.6, 128.6, 128.4, 127.9, 127.7, 127.5, 127.3, 126.5, 126.3, 125,3, 124.9, 124.4, 122.7, 39.1, 29.7 ppm. HR-MS: (ESI) calcd for $C_{16}H_{17}N_2$ $[M+H]^+$: 237.1392. Found: 237.1389.

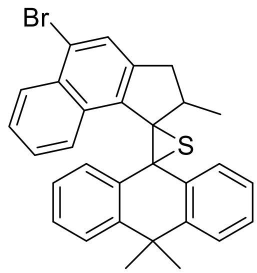

**120**

To an oven dried Schlenk flask charged with **GL-115** (0.708 g, 3.0 mmol) and $MgSO_4$ (200 mg), dichloromethane (20 mL) was added. The mixture was cooled down to −15 °C and $MnO_2$ (1.3 g, 15 mmol, Millipore-Sigma >90%) was added. The resulting mixture was stirred in a cold bath at ~−20 °C for 2 h. The solid was filtered. Thione **GL-119** (0.24 g, 1 mmol) was subsequentially added portion wise into the filtrate. The mixture was warmed to room temperature and stirred overnight. The solvents were removed by rotary evaporation. The crude product was purified by silica gel chromatography using hexane as the eluent, affording the title compound

**GL-120** in 51% yield (0.76 g). FTIR (KBr): 3062, 3027, 2972, 2922, 2864, 1602, 1580, 1559, 1508, 1463, 1445, 1385, 1360, 1323, 1281, 1256, 1213, 1156, 1046, 952, 919, 894, 793, 760, 712, 680 $cm^{-1}$. $^{1}H$ NMR (600 MHz, $CD_2Cl_2$) δ 9.47 (d, J = 8.38 Hz, 1H), 8.09-8.08 (m, 1H), 7.94-7.93 (m, 1H), 7.74-7.73 (m, 1H), 7.51-7.49 (m, 1H), 7.40 (s, 1H), 7.38-7.35 (m, 1H), 7.33-7.30 (m, 1H), 7.26-7.23 (m, 1H), 7.21-7.18 (m, 1H), 7.15-7.13 (m, 1H), 6.89-6.87 (m, 1H), 6.81-6.78 (m, 1H), 2.90 (m, 1H), 2.23 (d, J = 16.86 Hz, 1H), 1.97 (m, 1H), 1.66 (s, 3H), 1.10 (d, J = 7.15 Hz, 3H), 1.02 (s, 3H). $^{13}C$ NMR (150.8 MHz, $CD_2Cl_2$) δ 147.76, 147.55, 143.27, 137.81, 133.79, 132.77, 132.76, 131.88, 131.01, 127.96, 127.59, 127.39, 127.28, 126.72, 126.15, 125.97, 125.49, 125.02, 124.77, 124.21, 124.19, 123.89, 123.07, 73.74, 62.41, 39.36, 39.06, 37.21, 33.23, 26.51, 22.49.

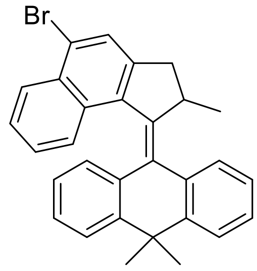


**GL-127**

To an oven dried Schlenk flask charged with **GL-120** (0.200 g, 0.40 mmol) and toluene (20 mL), 2 mL of hexamethylphosphoramide (HMPT) was added. The yellow solution was stirred and heated at 100 °C for overnight. The solution was permitted to cool to room temperature and the solvents were removed by rotary evaporation. The residues were purified by column chromatography on silica gel using hexanes/DCM (10/1) as the eluent, affording compound **GL-127** as a yellow solid (100 mg, 54%). FTIR (KBr): 3065, 3021, 2955, 2923, 2862, 1599, 1571, 1557, 1506, 1449, 1461, 1383, 1358, 1347, 1285, 1214, 1156, 1083, 952, 904, 890, 870, 759, 713, 579 $cm^{-1}$. $^{1}H$ NMR (600 MHz, $CDCl_3$) δ 8.07 (d, J = 8.37 Hz, 1H), 7.76-7.74 (m, 1H), 7.72 (s, 1H), 7.50-7.44 (m, 2H), 7.22-7.15 (m, 3H), 7.00-6.96 (m, 2H), 6.75-6.73 (m, 1H), 6.63-6.60 (m,

1H), 6.51-6.49 (m, 1H), 4.45 (m, 1H), 3.58 (dd, J1 = 15,24, J2 = 5.93 Hz, 1H), 2.53 (d, J = 15.24 Hz, 1H), 1.89 (s, 3H), 1.72 (s, 3H), 0.76 (d, J = 6.81 Hz, 3H). $^{13}$C NMR (150.8 MHz, $CDCl_3$) δ 147.21, 146.16, 145.56, 142.48, 140.09, 139.51, 136.35, 131.10, 130.21, 130.02, 128.28, 127.43, 127.37, 127.08, 126.98, 126.57, 126.27, 125.52, 125.36, 125.34, 124.89, 123.85, 123.38, 123.24, 40.54, 39.39, 37.73, 29.53, 24.68, 18.68. HRMS (ESI) calcd for $C_{30}H_{25}Br$ [M]: 464.1140. Found: 464.1133.

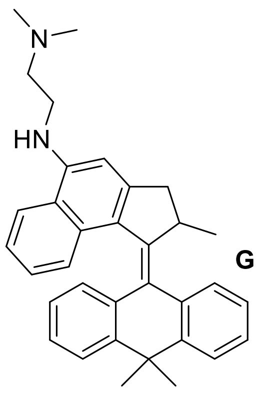

**GL-128**

(±)2,2′-Bis(diphenylphosphino)-1,1′-binaphthyl (BINAP) (15 mg, 0.024 mmol) and palladium(II) acetate (1.8 mg, 0.008 mmol) were mixed in toluene (5 mL). This solution was stirred for 20 min at room temperature. After that, NaO$^t$Bu (82 mg, 0.90 mmol), bromo-substituted motor GL-**127** (76 mg, 0.164 mmol) and *N*1,*N*1-dimethylethane-1,2-diamine (89 mg, 0.714 mmol) were added. The mixture was stirred and heated at 90 °C for overnight. The reaction mixture was permitted to cool to room temperature, and $H_2O$ (10 mL) and DCM (10 mL) were added. The organic phase was separated with a separation funnel. The aqueous phase was extracted with DCM (3 × 10 mL), and the organic phases were combined and washed with $H_2O$ (2 × 10 mL). After the solvents were removed by rotary evaporation, the residue was purified by column chromatography on silica gel using MeOH/DCM (1/9), obtaining **GL-128** as a yellow-green solid in 93% yield. FTIR (KBr): 3062, 3022, 2951, 2859, 2820, 2771, 1583, 1563, 1526, 1471, 1413, 1357, 1347, 1265, 1258, 1160, 1041, 761, 738, 711 cm$^{-1}$. $^1$H NMR (500 MHz, $CD_2Cl_2$) δ 7.89 (dd, J1 = 7.63,

J2 = 1.25 Hz, 1H), 7.76 (d, J = 8.61 Hz, 1H), 7.58-7.54 (m, 2H), 7.29-7.23 (m, 2H), 7.18 (t, 1H), 7.08-7.05 (m, 1H), 7.03 (d, J = 8.28 Hz, 1H), 6.79-6.77 (m, 2H), 6.63-6.61 (m, 2H), 4.48 (m, 1H), 3.63 (m, 1H), 3,36 (m, 2H), 2.76 (t, 2H), 2.35 (s, 6H), 0.85 (d, J = 6.77 Hz, 3H) ppm. $^{13}C$ NMR (125 MHz, $CD_2Cl_2$) δ 148.37, 147.21, 145.67, 145.49, 144.16, 141.27, 140.32, 129.77, 127.66, 127.30, 127.11, 126.92, 126.11, 125.73, 125.53, 125.48, 125.10, 124.95, 124.23, 124.01, 123.10, 122.99, 122.92, 122.78, 120.03, 101.71, 57.76, 57.54, 44.82, 40.86, 40.37, 39.97, 39.27, 37.84, 37.38, 29.69, 28.76, 24.47, 18.73, 18.34. HRMS (ESI) calcd for $C_{32}H_{37}N_2$ $[M+H]^+$: 473.2957. Found: 473.2952.

Synthesis of MM **3**

Final product characterization of MM **3** is reported in Supplementary Figs. 25-26.

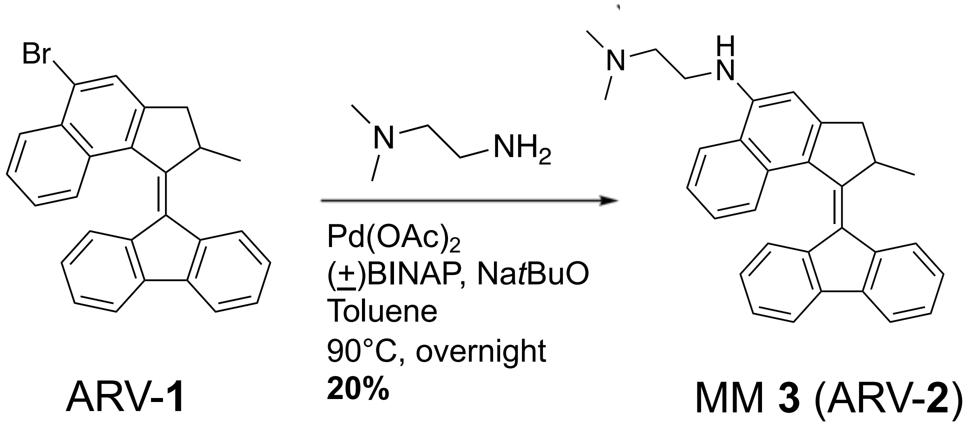


**Supplementary Scheme 2.** Synthesis of MM **3** (herein denoted ARV-**2**).

ARV-**1** was synthesized according to the previous literature[45]. ARV-**1** (247 mg, 0.58 mmol) and NaO*t*Bu (288 mg, 2.99 mmol) were added to an oven-dried screw-cap tube. The tube was evacuated and refilled with argon three times. To an oven-dried round bottom flask, (±) BINAP (50 mg, 15 mol %) and $Pd(OAc)_2$ (13 mg, 5 mol %) were added and the flask was evacuated and refilled with argon three times. Anhydrous toluene, degassed by bubbling with argon for 30 min, was added and the catalyst solution was stirred at room temperature for 30 min. At this time, the catalyst solution was transferred into the screwcap tube containing **13** and the NaO*t*Bu. *N,N*-dimethylethylenediamine (375 µL, 3.41 mmol) was added, the tube was capped, and the reaction was stirred at 90 °C overnight. Upon completion, the reaction was quenched with the addition aqueous ammonium chloride, extracted with DCM, and dried over sodium sulfate. The crude product was purified with column chromatography (2% MeOH in DCM) followed by preparatory TLC (2% MeOH in DCM) to afford **7** as a dark orange solid (50 mg, 20% yield).$^1$H NMR (500 MHz, DCM-d2) 8.00 (d, $J$= 7.7 Hz, 2H), 7.87 (dd, $J_1$= 7.4 Hz, $J_2$= 0.8 Hz, 1H), 7.81-7.79 (m, 1H), 7.66 (d, $J$= 8.3 Hz, 1H), 7.44-7.34 (m, 3H), 7.31-7.28 (m, 1H), 7.18 (td, $J_1$= 7.3 Hz, $J_2$= 0.9 Hz), 6.71 (d, $J$= 7.9 Hz, overlaps with signal at 6.69 ppm, 2H total), 6.69 (s, overlaps with signal at 6.71 ppm, 2H total), 6.00 (br s, 1H) 4.30-4.25 (m, 1H), 3.50 (dd, $J_1$=15.2 Hz, $J_2$= 5.7 Hz, 1H), 3.44-3.42 (m, 2H), 2.88-2.75 (m, 2H), 2.71 (d, $J$= 15.13 Hz, 1H), 2.39 (s, 6H), 1.40 (d, $J$= 6.9 Hz, 3H). $^{13}$C NMR (126 MHz, $CD_2Cl_2$) δ 153.39, 151.00, 147.01, 140.00, 139.01, 138.59, 137.40, 130.69, 127.93, 126.57, 126.51, 125.80, 125.58, 125.49, 125.47, 125.31, 124.43, 124.00, 123.48, 122.23, 121.04, 119.40, 118.71, 101.52, 57.28, 44.73, 44.69, 42.20, 40.53, 19.77. HRMS (ESI) *m/z* calculated for $[M+H^+]$ $C_{31}H_{30}N_2$ 431.2409; found, 431.2472.

Synthesis of MM **4**

Final product characterization of MM **4** is reported in Supplementary Figs. 25-26.

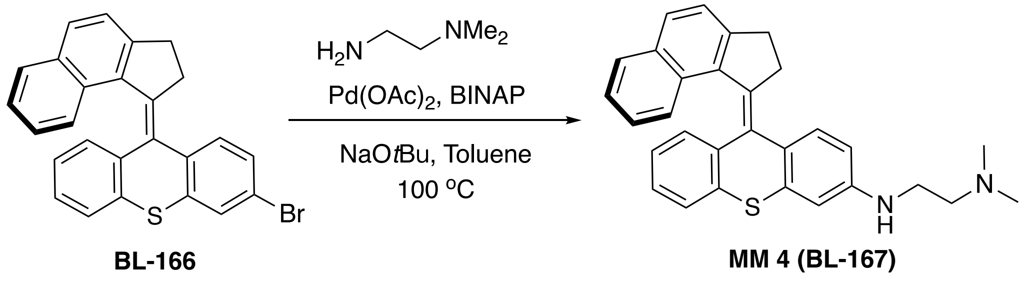


**Supplementary Scheme 3.** Synthesis of MM **4** (herein denoted **BL-167**).

**BL-166** was synthesized according to the previous literature [46]. BINAP (87 mg, 0.14 mmol) and palladium(II) acetate (15 mg, 0.07 mmol) were dissolved in dry toluene (15 mL). This solution was stirred for 30 min at room temperature, where upon it turned from dark red to dark orange. After this period NaO*t*Bu (456 mg, 4.75 mmol, 5 equiv) was added, followed by **BL-166** (420 mg, 0.95 mmol, 1 equiv) and $N^1$, $N^1$-dimethylethane-1,2-diamine (419 mg, 4.75 mmol, 5 equiv). The mixture was stirred at 100 °C for 2 d. Subsequently, the reaction mixture was poured into DCM (50 mL). After filtration, the solvents were evaporated. The crude product was dissolved in a small amount of DCM and purified using column chromatography (10% MeOH in DCM) to yield MM **4** as a yellow solid product (319 mg, 75% yield): $^1$H-NMR (600 MHz, $CD_2Cl_2$) δ 7.78 (d, *J* = 7.8 Hz, 0.5H), 7.75–7.62 (m, 2H), 7.61–7.52 (m, 1.5H), 7.47 (dd, *J* = 8.2, 2.9 Hz, 1H), 7.35 (t, *J* = 7.5 Hz, 0.5H), 7.24–7.09 (m, 2H), 7.01– 6.96 (m, 0.5H), 6.95–6.97 (m, 1H), 6.86–6.76 (m, 1.5H), 6.68–6.63 (m, 1H), 6.57 (t, *J* = 7.5 Hz, 0.5H), 6.46 (d, *J* = 8.4 Hz, 0.5H), 5.89 (dd, *J* = 8.4, 2.4 Hz, 0.5H), 4.80–4.00 (m, 2H), 3.25–3.17 (m, 4H), 3.05 (t, *J* = 5.9 Hz, 1H), 2.60 (t, *J* = 5.8 Hz, 1H),

2.50 (t, $J$ = 5.8 Hz, 1H), 2.27 (s, 3H), 2.22 (s, 3H); $^{13}$C NMR (151 MHz, $CD_2Cl_2$) δ 147.92, 147.84, 147.81, 147.38, 141.35, 139.99, 139.66, 139.63, 138.10, 137.82, 136.90, 136.69, 135.86, 135.67, 133.31, 133.26, 129.65, 129.58, 129.28, 129.18, 129.08, 128.75, 128.56, 128.48, 128.21, 128.11, 128.08, 128.06, 127.84, 127.51, 127.09, 126.64, 126.57, 126.38, 126.21, 126.12, 125.04, 124.99, 124.42, 124.39, 123.59, 123.57, 112.14, 112.07, 110.93, 110.90, 58.08, 57.96, 45.18, 45.13, 41.44, 41.31, 34.33, 34.30, 32.63, 32.59. ESI-MS: m/z 449.2 $[M+H]^+$; HRMS (ESI) calculated for [M+H, $C_{30}H_{29}N_2S]^+$: 449.1973, found: 449.2042. FTIR (cm$^{-1}$): 3052, 2938, 2852, 2819, 2762,1596, 1581, 1560, 1494, 1455, 1437, 1374, 1309, 1249, 1054, 1039, 956, 938, 812, 755, 743, 707.

### Supplementary Text

#### Excitation in a Non-Scanning Laser Regime

In typical experiments, the exciting laser for stimulation was rastered across a defined region-of-interest approximated as a circle of 5 µm diameter. We employed a second stimulation set-up on an epifluorescence microscope to verify that laser rastering is not necessary for excitation. Experiments with a non-rastering laser beam required similar stimulation powers to elicit responses from MM-treated cells (~$10^2$ W $cm^{-2}$).

We adapted the optical setup from a single cell fluorescence sensing optical system[47] to deliver a non-scanning laser beam (Coherent Chameleon Discovery) to the specimen from the top, at the excitation wavelength of the MM. The fluorescence signal is collected by an inverted fluorescence microscope from the bottom of the specimen, while an electronic shutter is used to release optical pulses for MM triggering at the desired frequency during fluorescence recording.

**Assessment of Stimulation Toxicity**

The purpose of this present work was to use MM to interact with cell signaling. However, several previous studies by our group have explored the same or similar molecules for inducing cell permeabilization and death[21,26-28,48]. These studies used minutes-long light doses rather than the millisecond-scale doses employed in this work. Nevertheless, assessing the effects of MM stimulation on cell viability remains imperative.

Potentially toxic effects of MM stimulation on cells were assessed in several ways. The results of these experiments are reported in Supplementary Figs. 7-9. First, qualitative analysis of stimulated cells was used to assess toxicity. For these experiments, HEK293 cells were treated with MM **1** and prepared as was typical for stimulation experiments (see *Methods*). At very high stimulation powers, cells exhibited membrane blebbing and runaway calcium accumulation (Supplementary Fig. 9a). By reducing the light power, we were able to find a more suitable light dose that elicited strong calcium responses from cells without observable toxicity (Supplementary Fig. 9b).

Second, once a suitable window of stimulation power had been identified, the effects of stimulation on cells were compared to healthy cells and several positive controls representing mechanisms of cell death. For these experiments, cells were loaded with Fluo-4, CellMask cell membrane labeling dye, and MM **1** and were imaged in the presence of cell-impermeant viability dye propidium iodide (PI; 1 μg $mL^{-1}$). The results of these experiments are shown in Supplementary Fig. 8. Apoptosis was initiated by prolonged exposure of cells to blue light, and the onset of apoptotic cell death in exposed colonies was confirmed by the observation of pyknosis (cell shrinkage)49, the uptake of small amounts of PI, and the enhanced fluorescence of Fluo-4 in the colony (Supplementary Fig. 7a). Necrotic cell death was observed by activating MM **1** *in situ* using a SOLA LED fed through a DAPI excitation filter (395/25 nm, 166 mW $cm^{-2}$) for 5 min (Supplementary Fig. 7b). Similar conditions have been shown to induce necrotic cell death in our previous work[28]. Using this protocol, cells do not shrink in the 5 min of stimulation, but they uptake large amounts of PI. Such changes were not observed in healthy, unperturbed cells (Supplementary Fig. 7c) or cells exhibiting MM-driven calcium responses driven by typical protocols (250 ms pulse to a 5 μm diameter at $3.2\times10^2$ W $cm^{-2}$ stimulation power; Supplementary Fig. 7d).

Finally, to account for potential aftereffects of stimulation, we grew isolated HEK293 colonies in a grid pattern by photolithographically patterning polydimethylsiloxane onto Ibidi imaging dishes. Colonies were seeded with 25,000 cells $cm^{-2}$. Two days after seeding, cells were treated with MM **1** and Fluo-4 using typical protocols (see *Methods*), and 10 selected cells were stimulated in selected colonies. Results from these experiments are shown in Supplementary Fig. 8. After 24 h, these colonies were imaged again in the presence of 1 μg $mL^{-1}$ PI to observe any effects of MM stimulation on growth, cell stress, or viability. The PI uptake of stimulated colonies was indistinguishable from untreated colonies. As a positive control, several colonies were irradiated with UV light (SOLA LED fed through a DAPI excitation filter (395/25 nm, 166 mW $cm^{-2}$) for 10 min on Day 3 of growth (MM **1** was not present). UV irradiation caused PI uptake into cells immediately and distinguished the UV-treated cells from cells that had been stimulated with MM **1** and light the day before.

In summary, qualitative assessment of the effects of MM stimulation on cells reveal that stimulation does not incur toxicity either by apoptosis or necrosis so long as the light dose administered is appropriately controlled.

**Effects of Rotation Speed and Directionality**

The propensity for unidirectional MM to rotate and exert mechanical force on their environment depend principally on two factors: the rotation rate and the photoconversion efficiency. Both of these depend on MM structure. The rotation rate is determined by the half-life of the thermal helix inversion, which is driven by the steric interactions governing the passage of the rotor over the stator[24]. The photoconversion efficiency, also known as the quantum yield of photoisomerization, represents the proportion of photons absorbed by the molecule that drive molecular rotation (as opposed to fluorescence or non-radiative decay processes). The quantum yield of photoisomerization is primarily influenced by the electronic structure. MM directionality can also be influenced by structure, as the driving force for unidirectional rotation depends upon the presence of a stereogenic center.

The selection of different MM of similar physicochemical character allowed us to probe the effects of altered MM speed or directionality on the signaling behavior elicited. MM **1,** which rotates at ~3 MHz, elicited the strongest signaling responses *in vitro*. ICW elicited by MM **1** peaked ~10-20 s after stimulation and decayed over the next minute. Slow-rotating MM **3** did not elicit any noticeable responses *in vitro*, whereas bidirectional MM **4** elicited inconsistent and weak responses. These results mirror those previously observed when tracking MM diffusion in solution (*20*) and when tracking their ability to permeabilize phospholipid bilayers[21]. In these experiments, slow-rotating MM were indistinguishable from solvent-only controls, whereas bidirectional MM were distinguishable from controls but did not exert as pronounced an effect as fast, unidirectional MM.

MM **2**, which has a reported rotation rate of ~43 MHz, also elicited strong responses *in vitro*, but did not elicit the same release kinetics, instead causing release of calcium to a stable level without decay until several minutes after stimulation. Given that ICW elicited by MM **1** reached a higher amplitude than those elicited by MM **2**, alongside the aforementioned difference in release and recovery kinetics, we conclude that MM **1** imparts a stronger stimulatory effect than MM **2** despite its slower rotation rate. The observed recovery by cells treated with MM **1** but not with MM **2** can be attributed to various mechanisms of cellular recovery, including SERCA activity and RyR closing probability, being enhanced by a larger cytosolic concentration of calcium *(1)*. If MM **2** does not elicit sufficient calcium release to cause accelerated cellular recovery, ICW elicited by MM **2** would persist at stable levels for a longer timeframe than those elicited by MM **1**. Since both these motors rotate in the MHz-regime, we anticipate that this difference in their activity can be attributed to their photoconversion efficiency. If a lower proportion of absorbed photons drive rotation in MM **2** as compared to MM **1**, this motor would impart a lower mechanical force than MM **1** despite its faster rotation rate. Indeed, previous studies of overcrowded alkene motor dynamics have shown that motors bearing similar core stator structures to MM **2** suffer from lower quantum yields compared to sulfur-bearing core stators[50].

In *Hydra*, several different trends are observed as compared to in cells. Most notably, MM **4** is able to drive non-trivial signaling behavior *in vivo*. MM **2** was also highly effective at driving *Hydra* contraction, and MM **1** and MM **2** were similar in ability to drive body column ICW. Small responses were observed even to stimulation with MM **3**, and the *Hydra* overall seemed a much more sensitive model system than HEK293 cells or cardiac myocytes. Compared to HEK293 cells, *Hydra* exhibit frequent spontaneous ICW and possess networks of cells that are excitable and electrically connected[35]. Hence, a response in one cell is more likely to be both amplified by a voltage-gated channel and propagated through gap junctions to other cells. The effectiveness of

MM **4** in *Hydra* but not in cultured cells illustrates how the smaller signals elicited by MM **4** can be amplified and propagated across cellular networks.

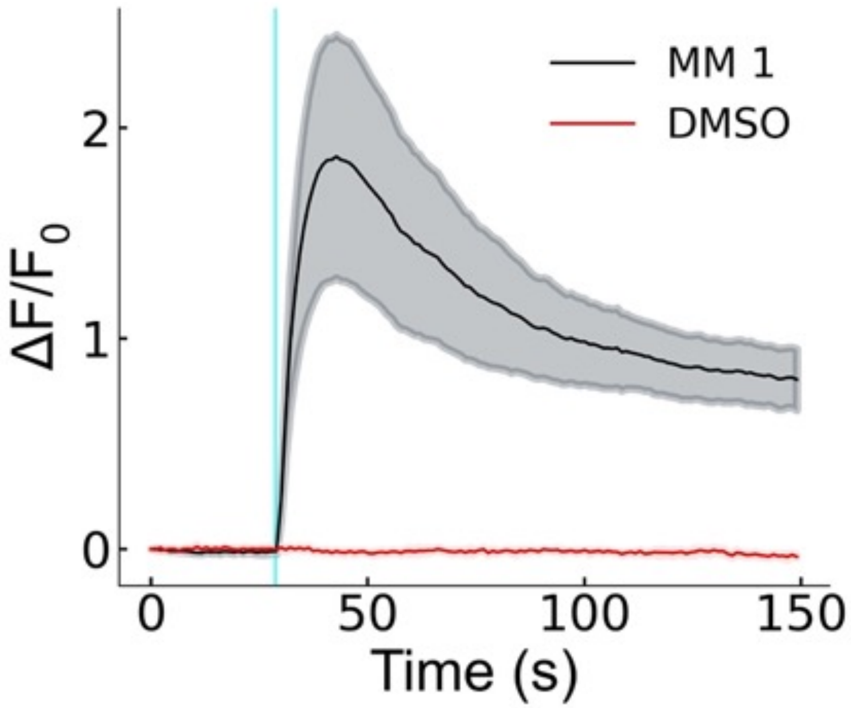


**Supplementary Fig. 1. Comparison to vehicle control.**

Normalized fluorescence intensity traces of Fluo-4 in HEK293 cells treated with either MM **1** (8 μM) or DMSO vehicle (0.1% v/v). The solid lines represent the average of n=6 cells. The shaded area represents the standard error of the mean.

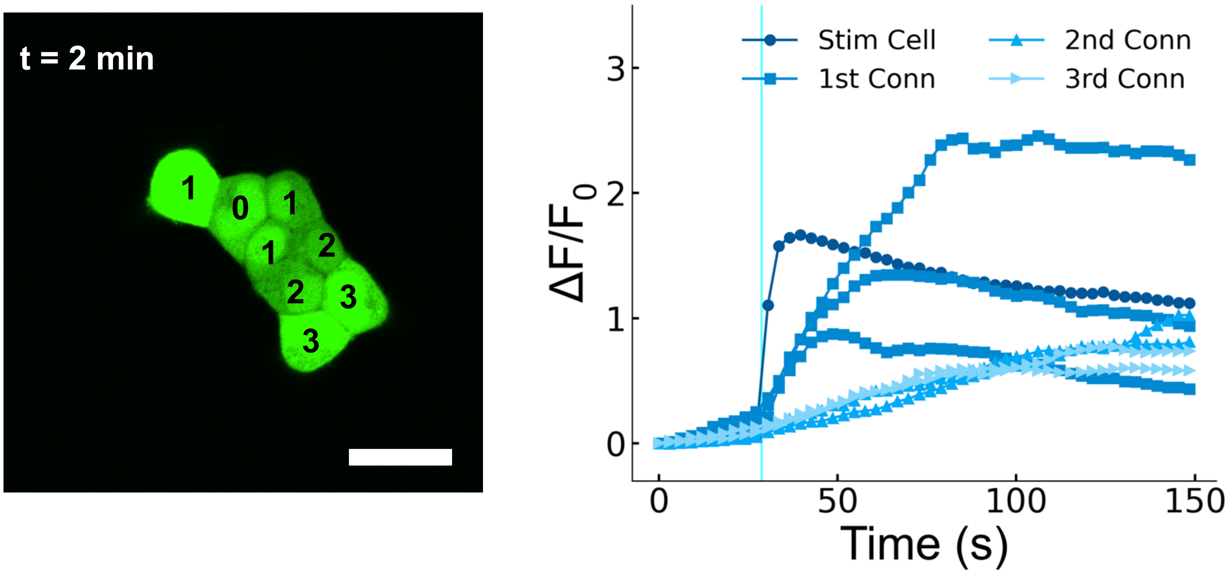


**Supplementary Fig. 2. MM-induced ICW propagate according to the degree of electrical connection in stimulated tissue.**

Fluo-4 fluorescence traces of HEK293 cells treated with MM **1** from a single colony (corresponding to Figure 1c), showing propagation of signal to surrounding cells over the timespan of several minutes. The cell marked "0" was stimulated with a 250 ms pulse of 400 nm light delivered to a 5 μm diameter circular region of interest at $3.2\times10^{2}$ W $cm^{-2}$. The image on the left shows the cells from which the traces are derived. The numbers on each individual cell denote the separation, in number of cells, from the stimulated cell. The plot on the right shows normalized intensity traces of both the stimulated cell and the other cells in the colony. The cells marked "$1^{st}$ Conn" were directly connected to the stimulated cell and correspond to the cells marked "1" in the left image. The cells marked "$2^{nd}$ Conn" were connected to a cell designated "$1^{st}$ Conn" and correspond to cells marked "2" in the left image. The cell marked "$3^{rd}$ Conn" were connected to cells designated "$2^{nd}$ Conn" and correspond to the cells marked "3" in the left image. Scale bar is 20 μm.

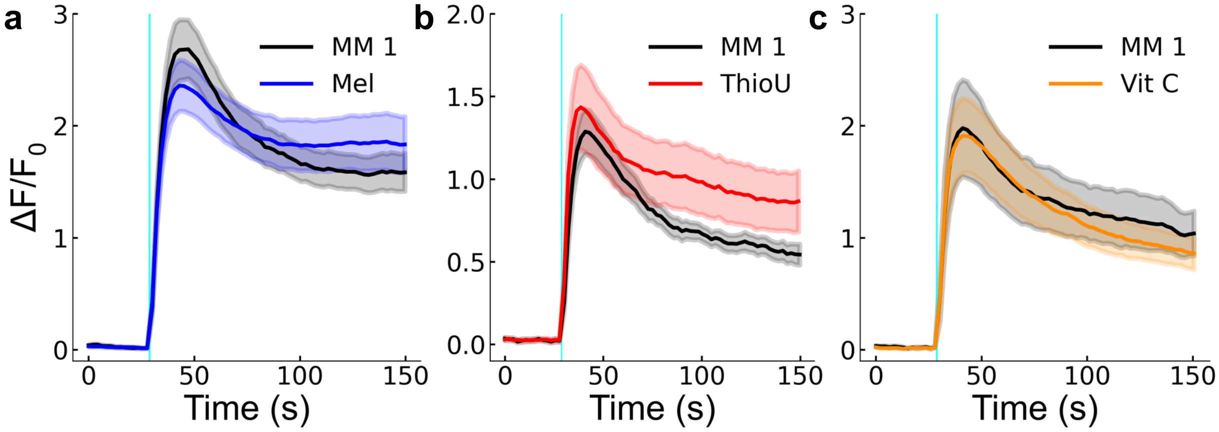


**Supplementary Fig. 3. Administration of ROS scavengers does not affect MM induced ICW.** Normalized fluorescence intensity traces of HEK293 cells treated with MM **1** and Fluo-4 and administered **(a)** melatonin (Mel; 100 µM), **(b)** thiourea (ThioU; 50 mM), and **(c)** L-ascorbic acid (Vit C; 2 mM). The shaded region indicates the standard error of n=6 stimulation attempts. Dark traces indicate positive controls collected prior to treatment. Stimulation was administered at $4.5\times10^{2}$ W $cm^{-2}$. Cells were incubated with chosen ROS scavengers for 1 h prior to stimulation.

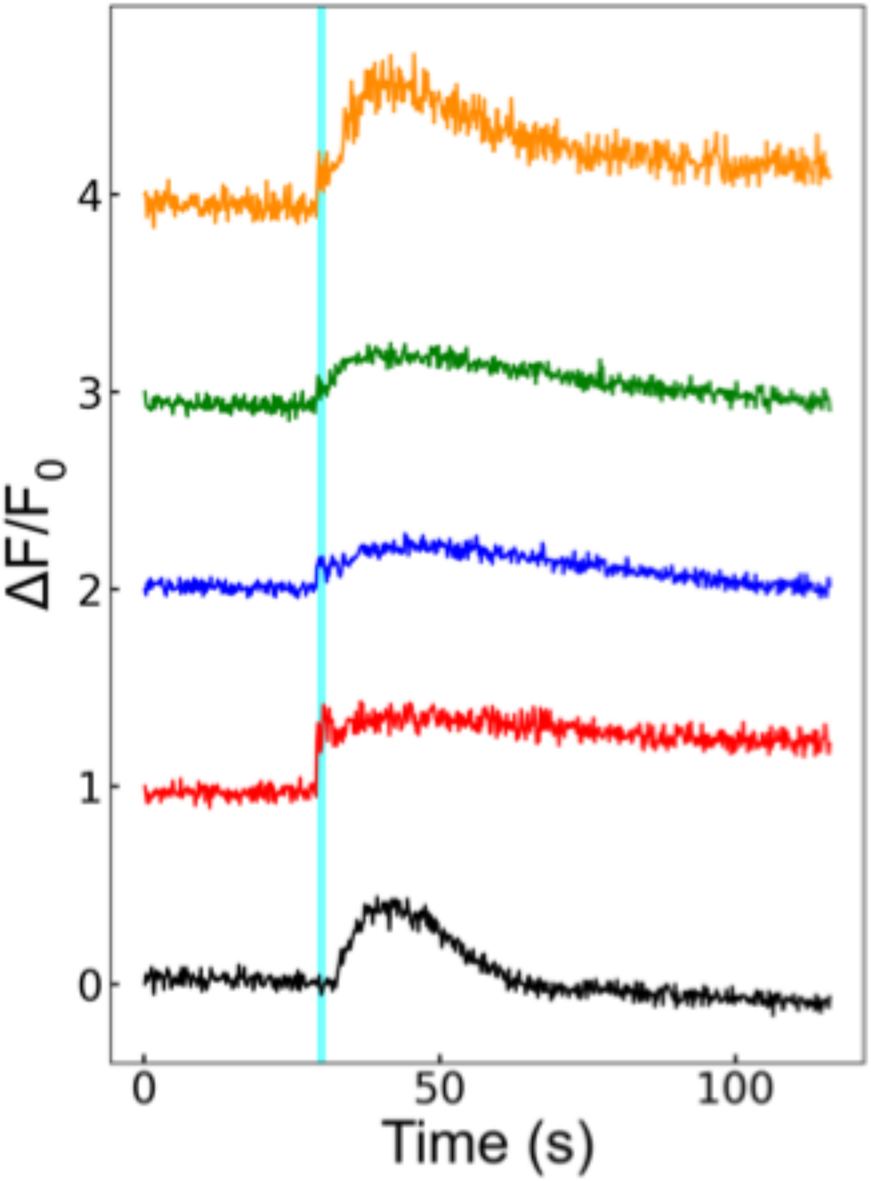


**Supplementary Fig. 4. Using an alternative calcium tracking dye.**

Normalized fluorescence intensity traces of X-Rhod-1 in HEK293 cells treated with MM **1** and stimulated with a 250 ms pulse of 400 nm light delivered to a 5 µm diameter circular region of interest at $3.2\times10^{2}$ W cm$^{-2}$. Different colors represent distinct cells. X-Rhod-1 was loaded into cells at 2 µM over a period of 45 min and excited using 561 nm laser light in a Nikon A1 Rsi fluorescence microscope.

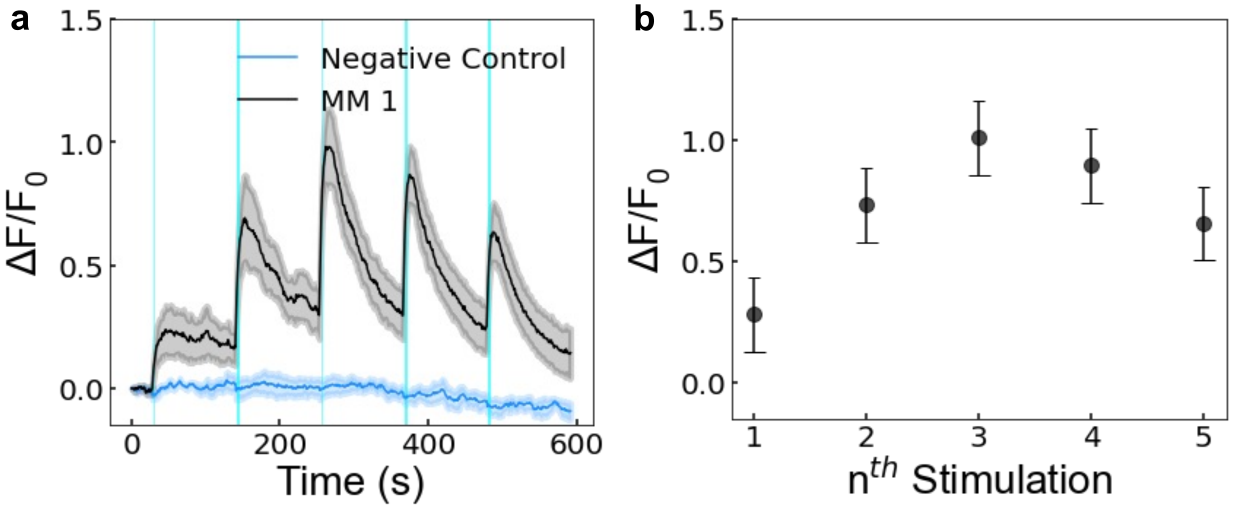


**Supplementary Fig. 5. Calcium responses to MM are repeatable.**

**(a)** Fluorescence traces of Fluo-4 in HEK293 cells treated with MM **1** as described in the Methods section and stimulated multiple times ($3.2\times10^{2}$ W $cm^{-2}$, 5 µm diameter, 250 ms pulse time). The "negative control" represents vehicle-treated (0.1% DMSO) cells that were subjected to the same irradiation regime as MM-treated cells. The shaded region represents the standard error of the mean across n=6 cells. **(b)** Magnitude of changes in intracellular calcium levels in MM **1**-treated cells exposed to multiple light stimulations. Error bars represent the standard error of n=6 cells. The cyan line indicates the time of stimulus presentation.

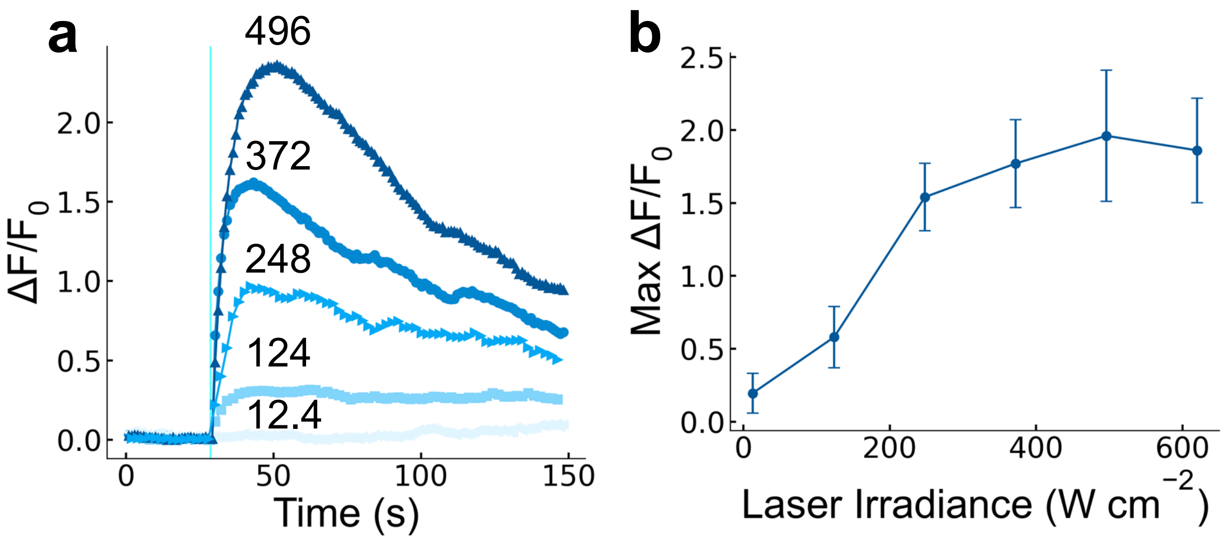


**Supplementary Fig. 6. Light power dependence of the responses of HEK293 cells to MM 1.**

**(a)** Representative fluorescent intensity traces of Fluo-4 over time at varying laser duty cycles, showing higher response amplitude at higher irradiance. Numbers in (a) depict the irradiance of each experiment in W cm$^{-2}$. Stimuli were delivered as a 250 ms pulse to a 5 µm diameter circular area. **(b)** Dose-response curve showing higher calcium response amplitudes at higher light intensities. Values shown represent the peak response amplitude. Error bars represent the standard error of the mean of at least six individual cells.

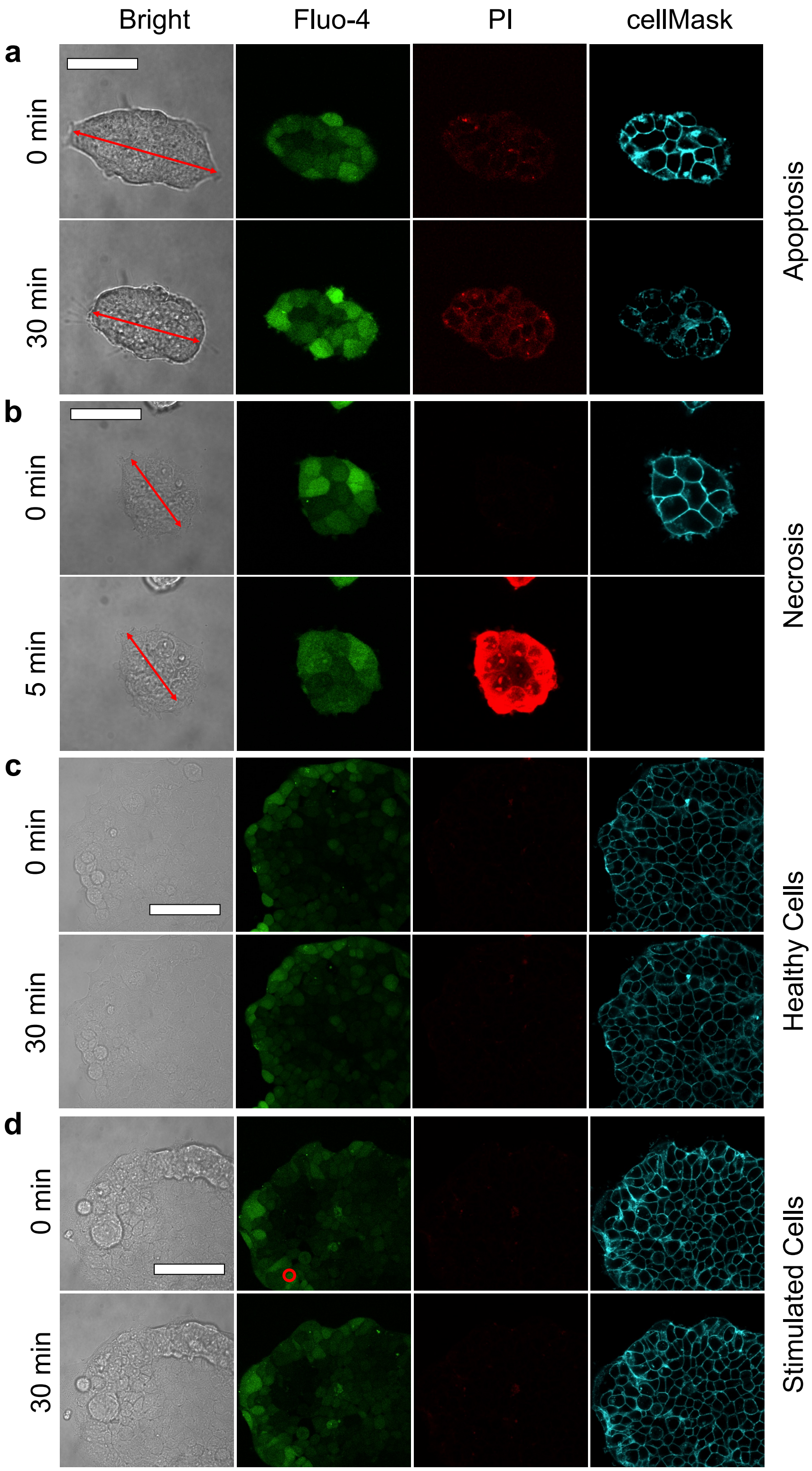
Bright
Fluo-4
PI
cellMask
a
0 min
30 min
Apoptosis
b
0 min
5 min
Necrosis
c
0 min
30 min
Healthy Cells
d
0 min
30 min
Stimulated Cells

**Supplementary Fig. 7. MM-induced ICW do not cause apoptosis or necrosis.**

Cells were treated with MM **1** (8 µM) as described in Methods and stained with Fluo-4 (green) to track calcium flux, propidium iodide (PI; red) to track membrane poration and cell death, and cellMask plasma membrane stain (cyan) to distinguish and track individual cells. **(a)** Positive control for apoptotic cell death with characteristic cell shrinkage (pyknosis) incurred by consecutive cell exposure to blue light (488 nm, irradiance of ~60 W $cm^{-2}$ rastered across the entire image for a period of 30 min). **(b)** Positive control for necrotic cell death incurred by treatment of cells with MM **1** and irradiation with UV light (365-385 nm, ~0.16 W $cm^{-2}$ for 5 min). Cell death is indicated by rapid PI uptake. **(c)** Negative control showing morphology of undisturbed, healthy cells. **(d)** A small colony of cells shown just before and 30 min after a typical stimulation experiment with the site of stimulation marked (red circle, second image on top row). All scale bars are 50 µm and apply to the images in their sub-figure.

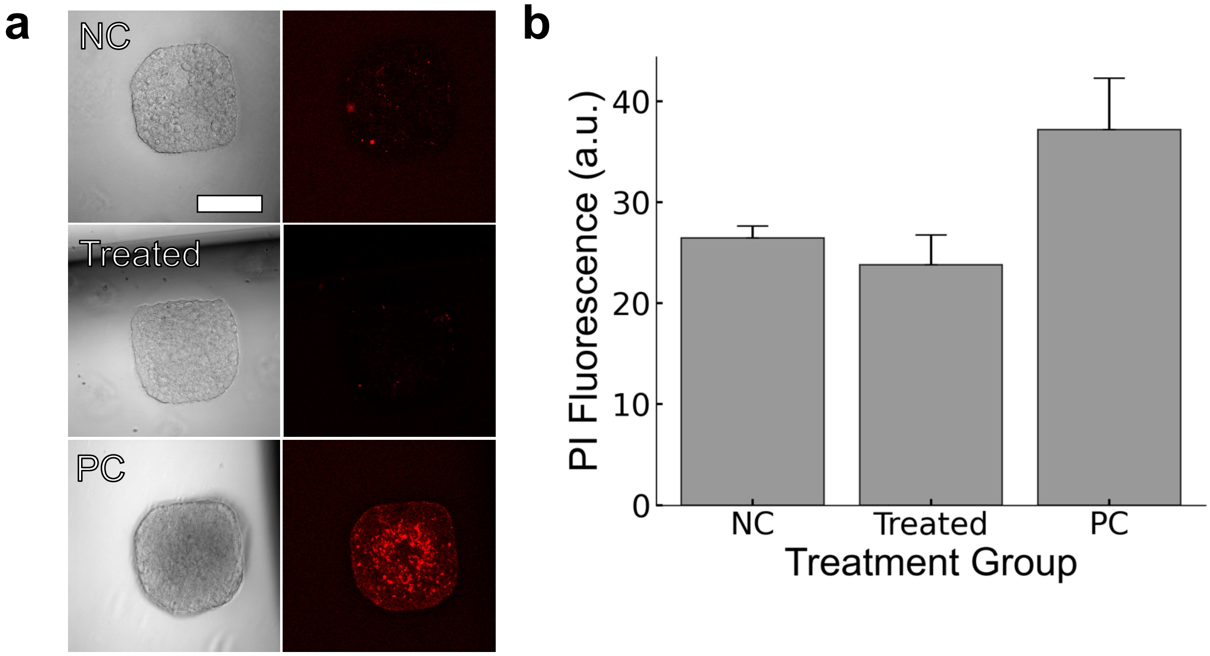


**Supplementary Fig. 8. MM-induced ICW do not have toxic effects on stimulated cell populations.**

Ibidi microscope dishes were photolithographically patterned with polydimethylsiloxane to prevent cell attachment except for in a gridded array pattern, resulting in isolated HEK293-colonies of defined size and geometry[51]. These colonies were imaged 3 days after being seeded following i) no treatment (negative control; NC), ii) stimulation with MM **1** and light on day 2 (10 single-cell stimulations per colony, $3.2\times10^{2}$ W cm$^{-2}$; Treated) , or iii) irradiation with UV light (365-385 nm, ~0.16 W cm$^{-2}$) for 5 min on day 3 (positive control; PC). **(a)** Bright field (left) and fluorescent (right) images of representative colonies from each treatment showing the onset of PI uptake into colonies in the PC group and no substantial uptake post stimulation in the treated cells. Scale bar is 200 µm and applies to all images. **(b)** Calculated PI fluorescent intensity. Error bars indicate standard deviation. n=3 colonies for each group.

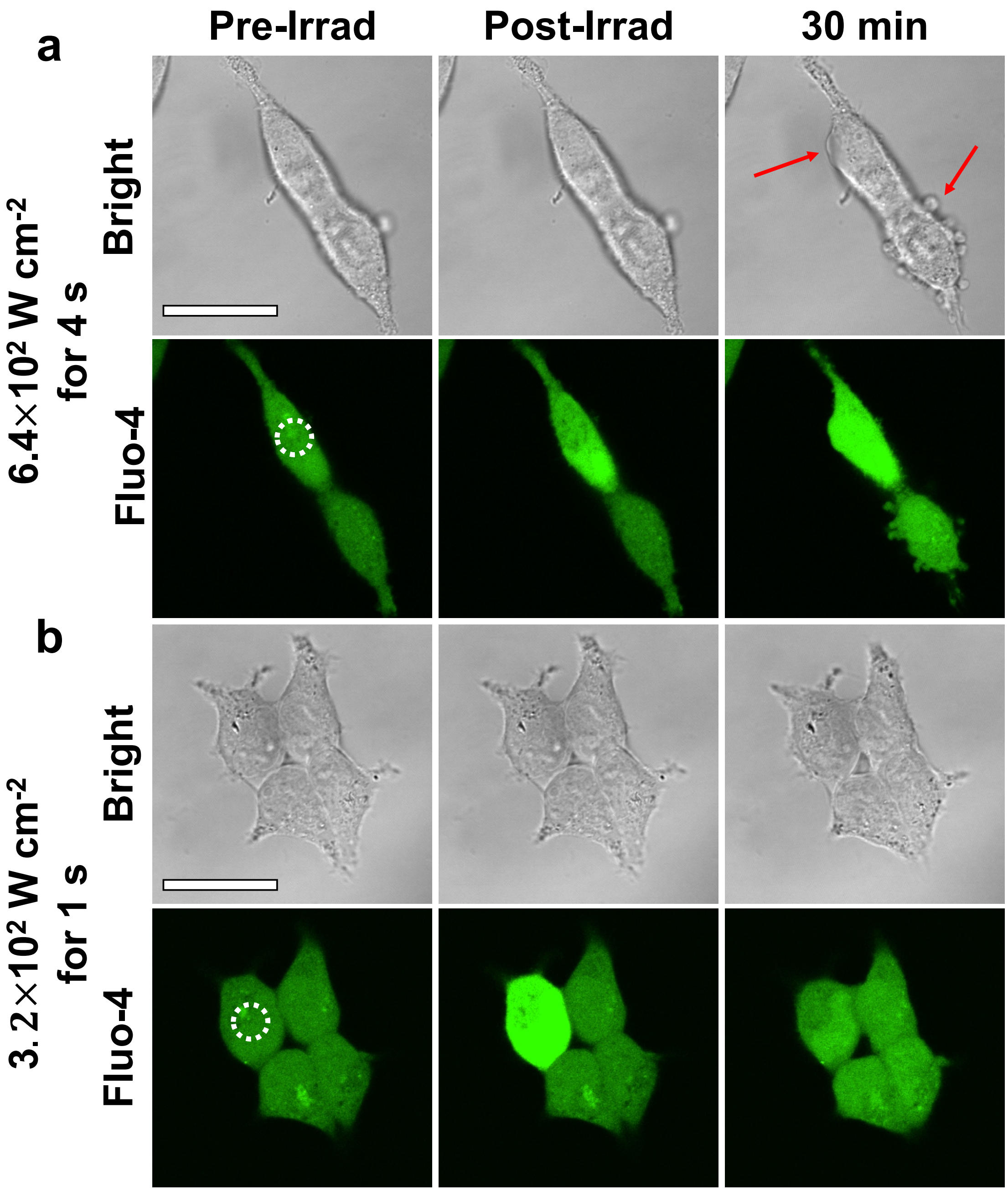


**Supplementary Fig. 9. Toxicity at high exposure times and high laser power.**

(**a**) Confocal microscope images of HEK293 cells treated with MM **1** and Fluo-4 calcium tracking dye after a 4 s light pulse of 400 nm light at $6.4\times10^2$ W cm$^{-2}$. Thirty minutes after treatment, the calcium signal has not returned to homeostasis, and membrane blebbing is apparent (red arrows). **(b)** Confocal images of HEK293 cells treated with MM **1** and Fluo-4 calcium tracking dye after laser pulses of $3.2\times10^2$ W $cm^{-2}$ for 1 s. After 30 min, the calcium signal has returned to homeostatic levels, and the stimulated cell and surrounding cells display no apparent damage. Scale bars are 50 μm and apply to all images in their sub-figure. The white dotted circles indicate the area of stimulation.

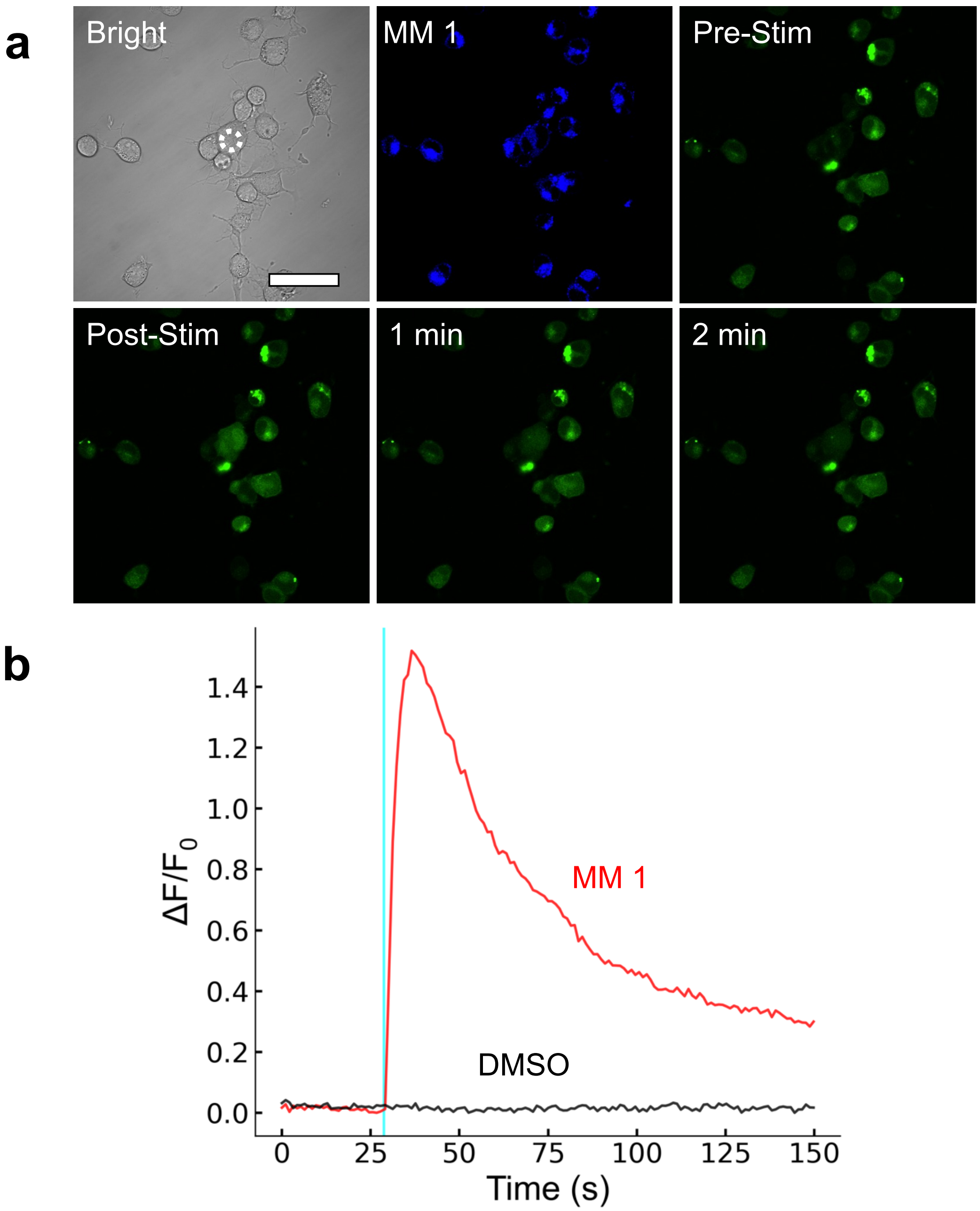


**Supplementary Fig. 10. ICW in N2A cells after stimulation with MM 1 and light.**

**(a)** Confocal images of N2A cells treated with MM **1** (blue) and calcium tracking dye (Fluo-4; green). The white circle in the top-left image depicts the area to which stimulus was delivered. The scale bar is 50 µm and applies to all images. **(b)** Fluo-4 fluorescence traces of N2A cells treated with MM **1** and stimulated with 250 ms pulse of 400 nm light delivered to a 5 μm diameter circular region of interest at $3.2\times10^{2}$ W $cm^{-2}$ stimulation power.

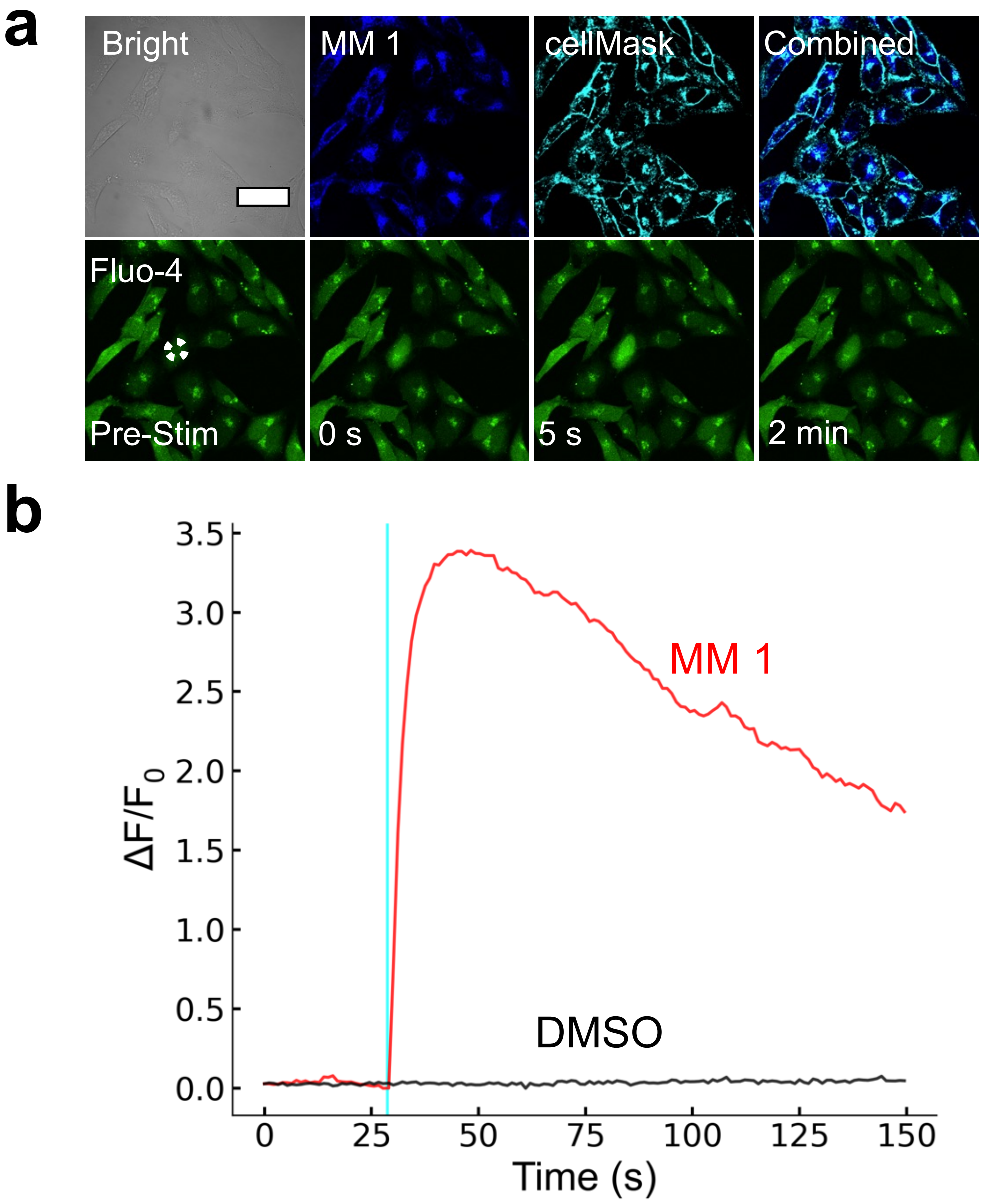


**Supplementary Fig. 11. ICW in HeLa cells after stimulation with MM 1 and light.**

**(a)** Confocal images of HeLa cells treated with MM **1** (blue) and Fluo-4 calcium tracking dye (green). The white circle in the bottom-left image depicts the area to which light stimulus was delivered. Scale bar is 50 µm and applies to all images. **(b)** Fluo-4 fluorescence traces of HeLa cells treated with MM **1** and stimulated with 250 ms pulse of 400 nm light delivered to a 5 μm diameter circular region of interest at $3.2\times10^{2}$ W $cm^{-2}$ stimulation power.

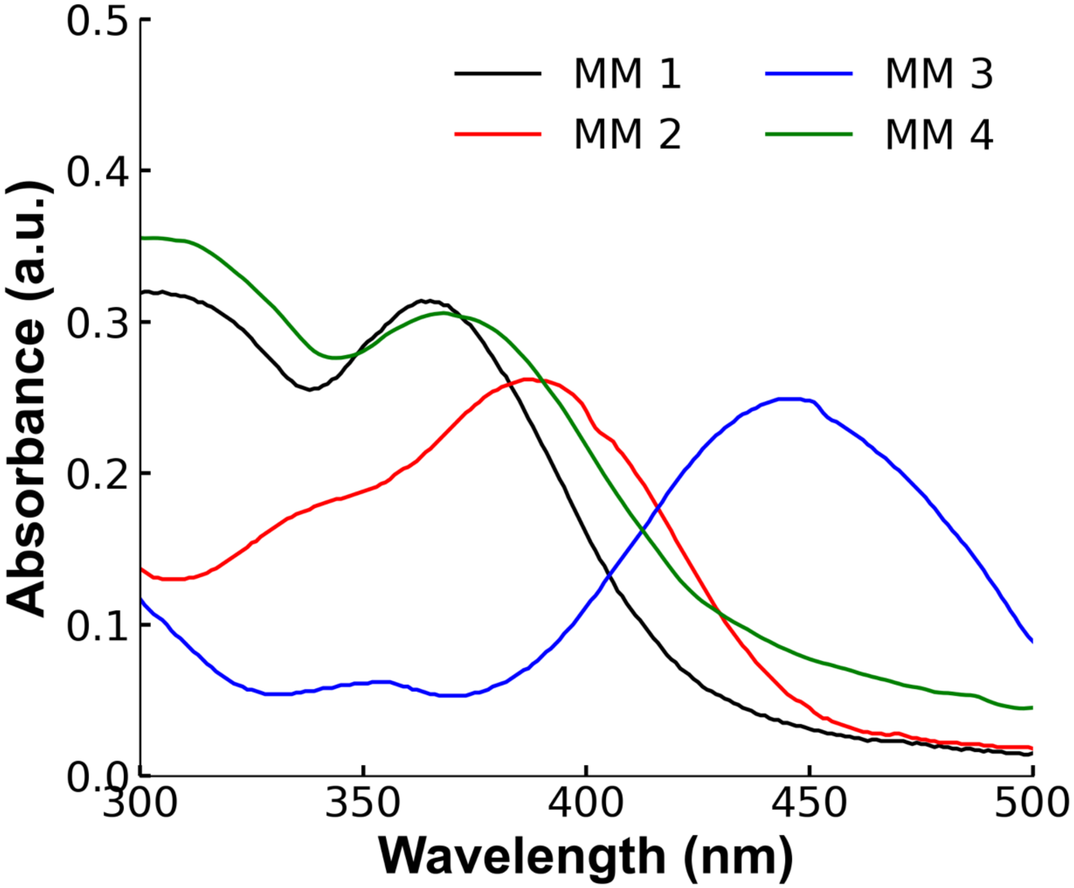


**Supplementary Fig. 12. UV-Vis spectra.**

Sample UV-Vis spectra of each MM taken at 16 μM in spectral-grade water.

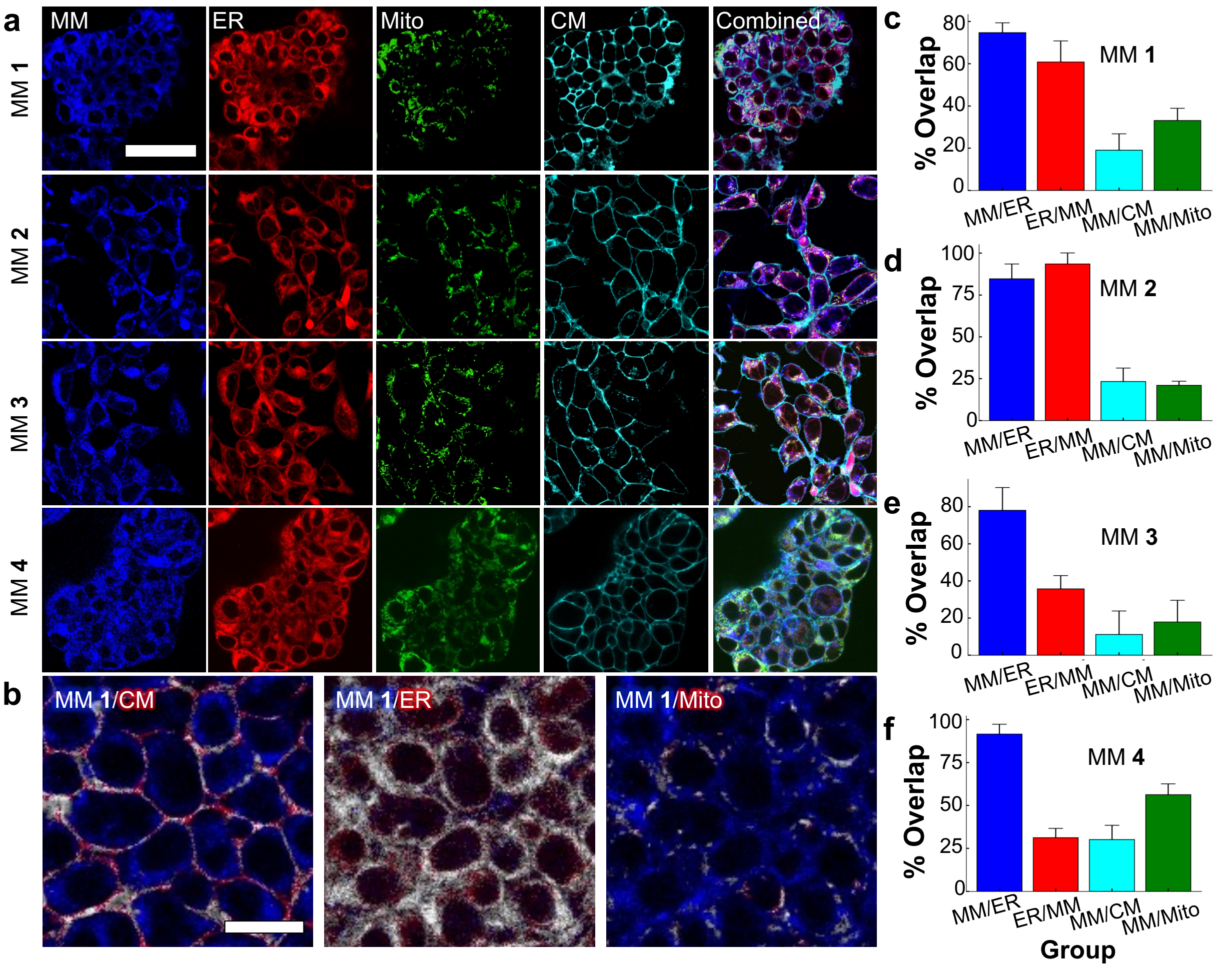


**Supplementary Fig. 13. Intracellular location of each MM.**

**(a)** Confocal images of HEK293 cells incubated with each MM, and different organelle-targeting dyes, namely ER-Tracker Red (ER), MitoTracker Green (mitochondria), and cellMask (plasma membrane), showing that MM preferentially distribute to the ER. The 50 μm scale bar applies to all images in the first four rows. **(b)** Pixel overlap maps of MM **1** fluorescence paired with fluorescence from organelle-targeted dyes. Blue color indicates pixels with fluorescence from MM **1** but not dye. Red color indicates pixels with fluorescence from dye but not MM **1**. Gray color indicates pixels with overlapping fluorescence from both MM **1** and dye. Left: cell membrane; middle: ER; right: mitochondria. Scale bar is 20 μm and applies to all images. Percent volumetric coverage of each organelle by **(c)** MM **1**, (**d)** MM **2**, **(e)** MM **3**, and (**f**) MM **4**. MM/ER depicts the percentage of MM found in the ER, while ER/MM depicts the percentage of ER covered with MM. MM/CM and MM/Mito depict the proportion of MM found in the cell membrane and mitochondria, respectively.

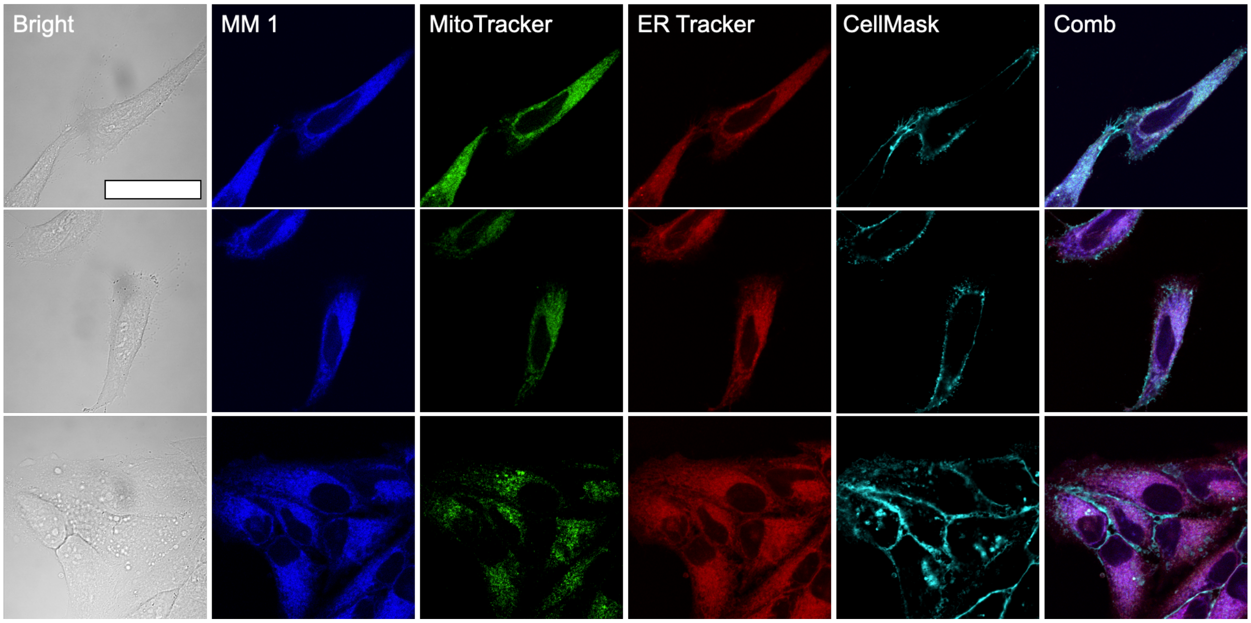


**Supplementary Fig. 14. Location of MM in HeLa cells.**

Microscopy images revealing the intracellular localization of MM **1** (blue), MitoTracker Green (green), ER-Tracker-Red (red), and cellMask plasma membrane dye (cyan) in HeLa cells. Scale bar is 50 μm and applies to all images.

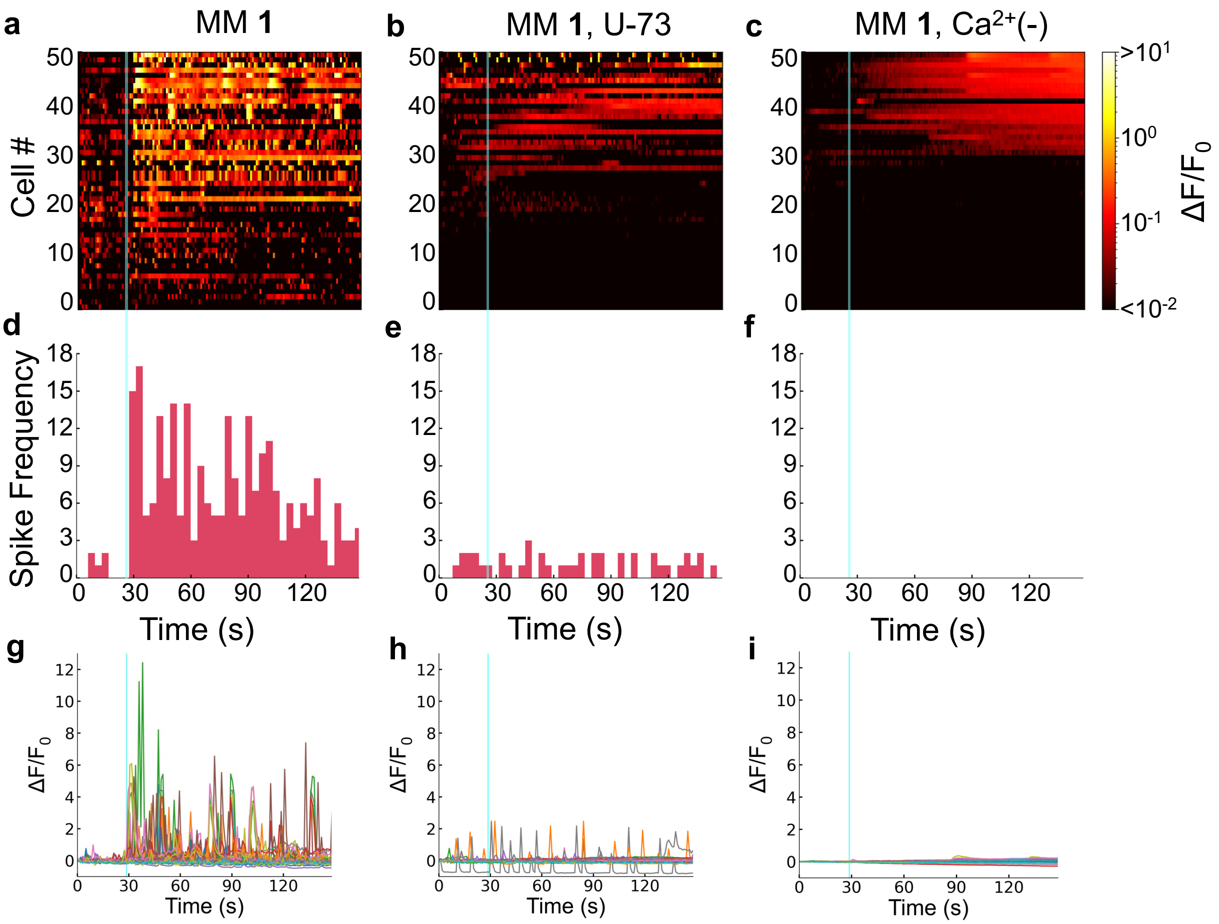


**Supplementary Fig. 15. Analysis of signaling activity of cardiomyocytes in colonies connected to cells stimulated by MM 1.**

Heatmap plots **(a-c),** temporal spike frequencies **(d-f)**, and individually plotted traces **(g-i)** of cardiomyocytes adjacent to a cell stimulated with **(a,d,g)** MM **1** + light, **(b,e,h)** MM **1** + light in 10 μM U-73, and **(c,f,i)** MM **1** + light in PBS. Light stimulation was performed using a 5.1×10$^2$ W cm$^{-2}$ 250 ms pulse of 400 nm laser light delivered to a 5 μm diameter area. The histograms in (d-f) indicate the time distributions of calcium spikes in cardiomyocytes identified by a peak detection algorithm (see Methods). Different colors in the individual traces in (g-f) indicate individual cells.

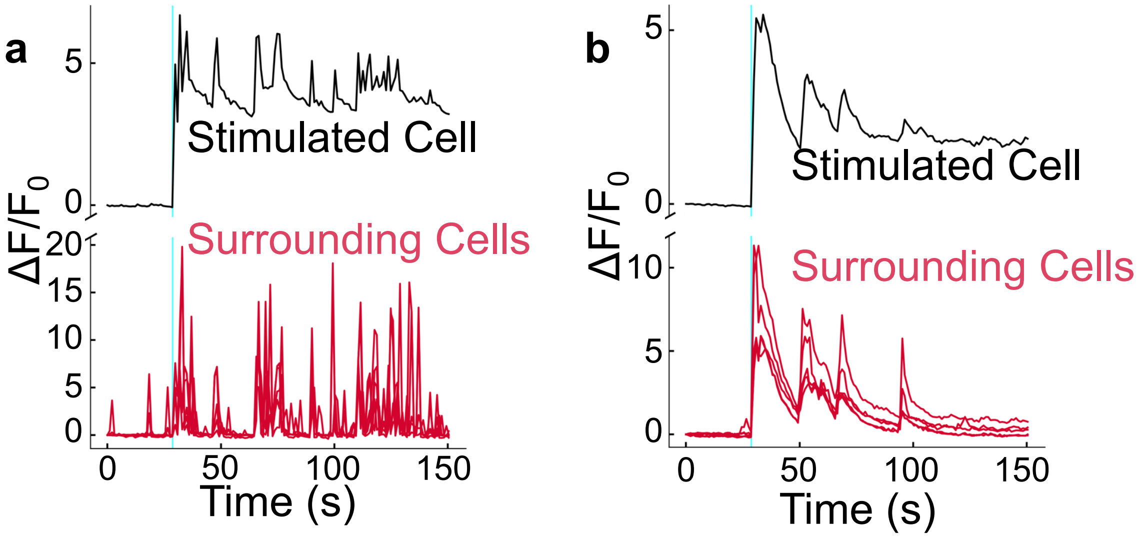


**Supplementary Fig. 16. Different phenotypes of cardiac myocyte activation in surrounding cells.**

Response phenotypes of individual colonies of cardiac myocytes showing (**a**) enhanced firing due to membrane depolarization and (**b**) synchronous calcium increase. Black traces indicate the stimulated cell. Red traces indicate adjacent cells in a colony. Light stimulation was performed using a $5.1 \times 10^2$ W $cm^{-2}$ 250 ms pulse of 400 nm laser light delivered to a 5 μm diameter area.

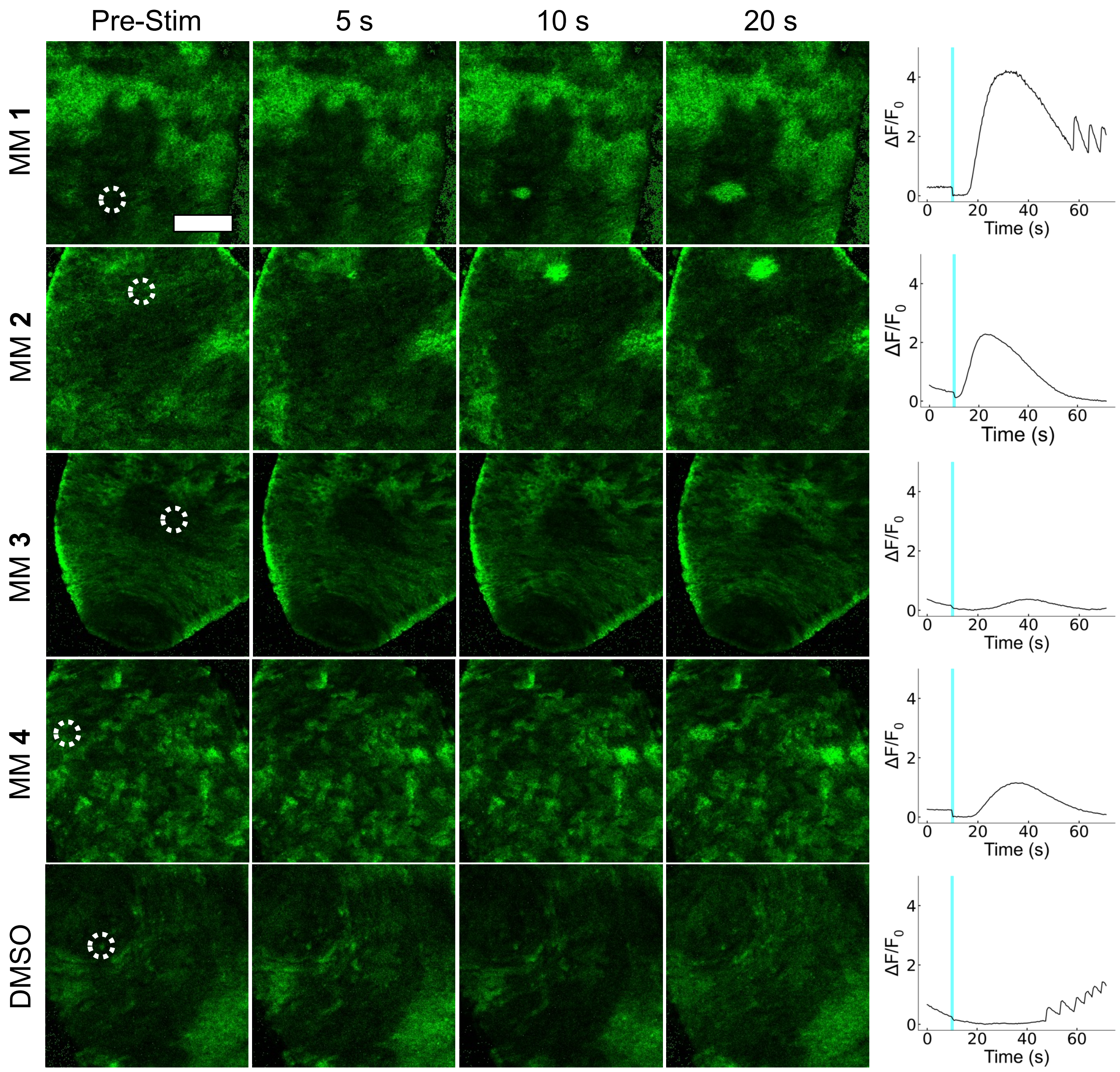


**Supplementary Fig. 17. Representative results for regional ICW elicited in MM-treated *Hydra vulgaris.***

Representative images (left) and normalized fluorescence intensity traces of GCaMP7b (right) of *Hydra* treated with each MM (24 μM) or DMSO (0.3% v/v). Scale bar is 100 μm and applies to all images. Traces were taken from the area of stimulation shown by the white dotted circles.

*Hydra* were stimulated at $9.0\times10^2$ W $cm^{-2}$ power with a 405 nm laser in a 10 µm diameter circular area for 1 s (Protocol **I**).

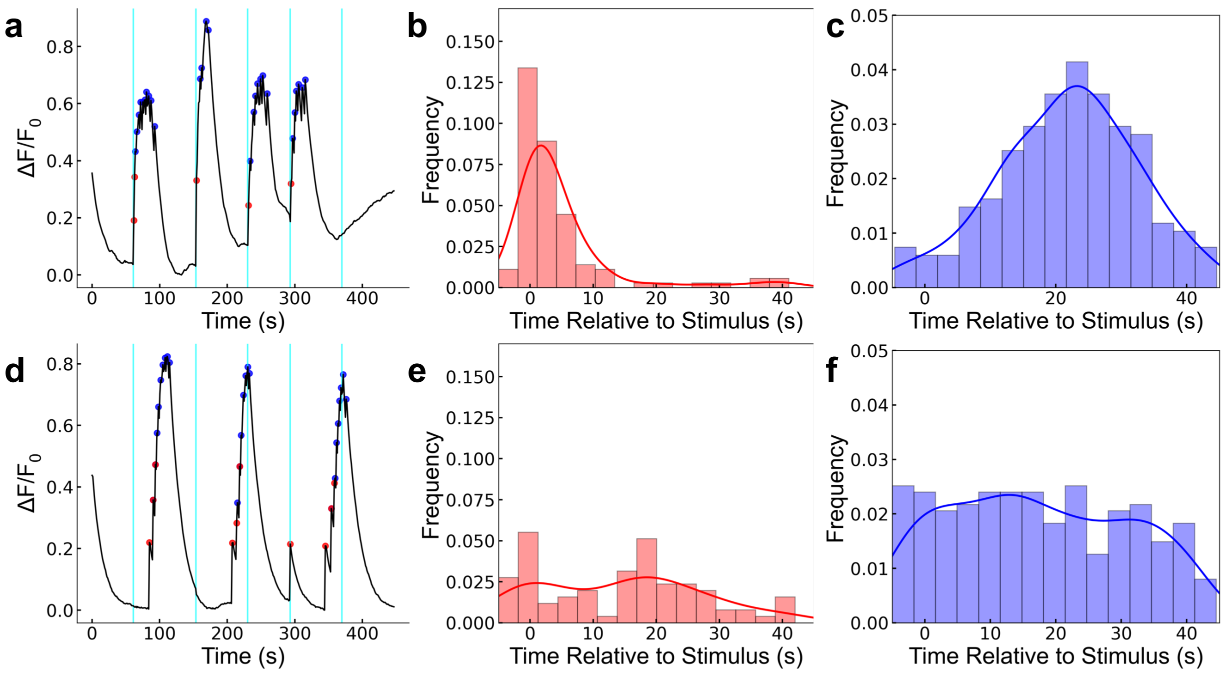


**Supplementary Fig. 18. Peak analysis of MM 1-treated *Hydra* (rotation rate of 3 MHz).**

**(a)** Characteristic GCaMP7b fluorescence trace of *Hydra* treated with MM **1** + light (Protocol **II**). , The contractions (peaks) most often occur in concert with stimuli (cyan lines). **(b)** Temporal location of contraction onset relative to presentation of light stimulus. **(c)** Temporal location of contraction burst peaks relative to presentation of light stimulus. **(d)** Characteristic GCaMP7b fluorescence trace of *Hydra* treated with MM **1** and sham light stimulus (same protocol with 0% laser duty cycle). Contractions do not occur in concert with sham stimulus. **(e)** Temporal location of contraction onset peaks relative to presentation of sham light stimulus. **(f)** Temporal location of contraction burst peaks relative to presentation of sham light stimulus. Histograms were calculated across all collected data (at least 50 stimulation or sham stimulation attempts in at least 5 *Hydra*).

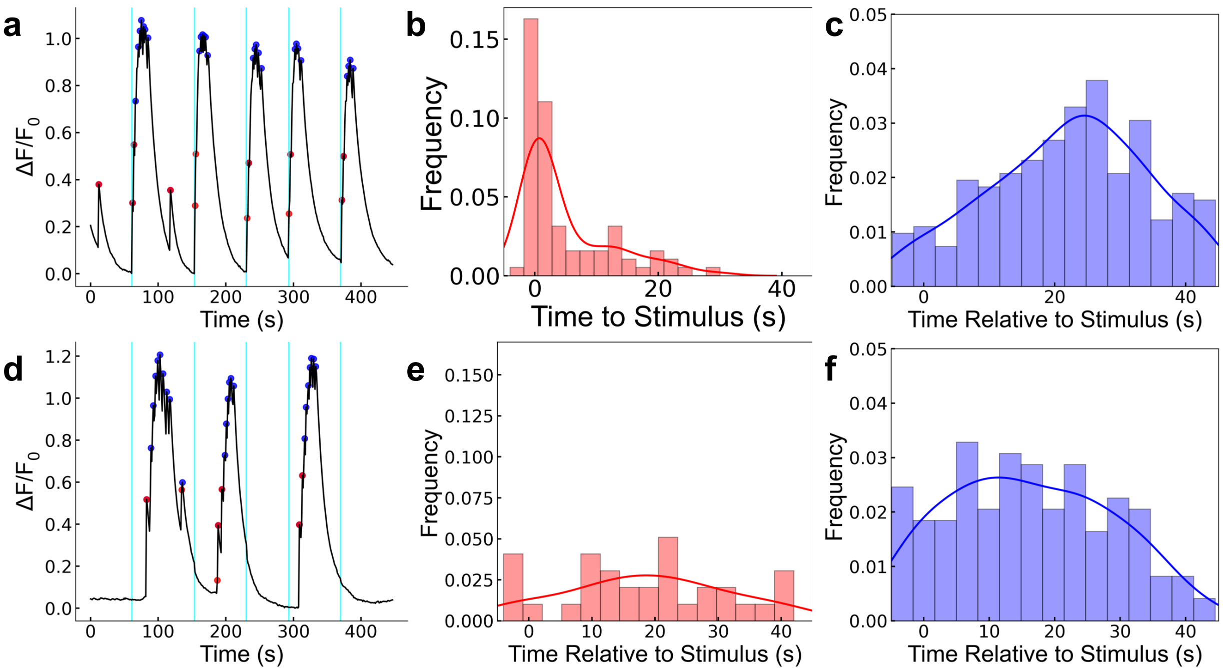


**Supplementary Fig. 19. Peak analysis of MM 2-treated *Hydra* (fastest MM; rotation rate of 43 MHz).**

**(a)** Characteristic GCaMP7b fluorescence trace of *Hydra* treated with MM **2** + light (Protocol **II**). The contractions (peaks) occur in concert with stimuli (cyan lines). **(b)** Temporal location of contraction onset relative to presentation of light stimulus. **(c)** Temporal location of contraction burst peaks relative to presentation of light stimulus. **(d)** Characteristic GCaMP7b fluorescence trace of *Hydra* treated with MM **2** and sham light stimulus (same protocol with 0% laser duty cycle). Contractions do not occur in concert with sham stimulus. **(e)** Temporal location of contraction onset peaks relative to presentation of sham light stimulus. **(f)** Temporal location of contraction burst peaks relative to presentation of sham light stimulus. Histograms were calculated across all collected data (at least 50 stimulation or sham stimulation attempts in at least 5 *Hydra*).

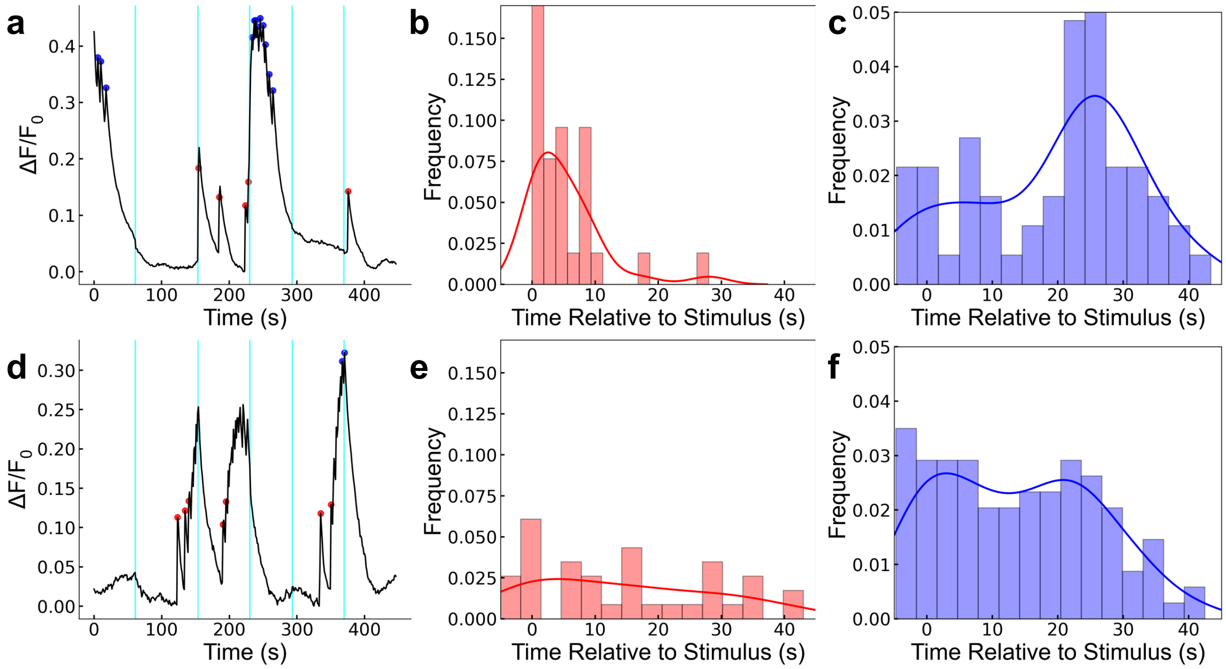


**Supplementary Fig. 20. Peak analysis of MM 3-treated *Hydra* (slow MM; rotation rate of $10^{-1}$ Hz).**

**(a)** Characteristic GCaMP7b fluorescence trace of *Hydra* treated with MM **3** + light (Protocol **II**). The contractions (peaks) occasionally follow stimuli (cyan lines). **(b)** Temporal location of contraction onset relative to presentation of light stimulus. **(c)** Temporal location of contraction burst peaks relative to presentation of light stimulus. **(d)** Characteristic GCaMP7b fluorescence trace of *Hydra* treated with MM **3** and sham light stimulus (same protocol with 0% laser duty cycle). Contractions do not occur in concert with sham stimulus. **(e)** Temporal location of contraction onset peaks relative to presentation of sham light stimulus. **(f)** Temporal location of contraction burst peaks relative to presentation of sham light stimulus. Histograms were calculated across all collected data (at least 50 stimulation or sham stimulation attempts in at least 5 *Hydra*).

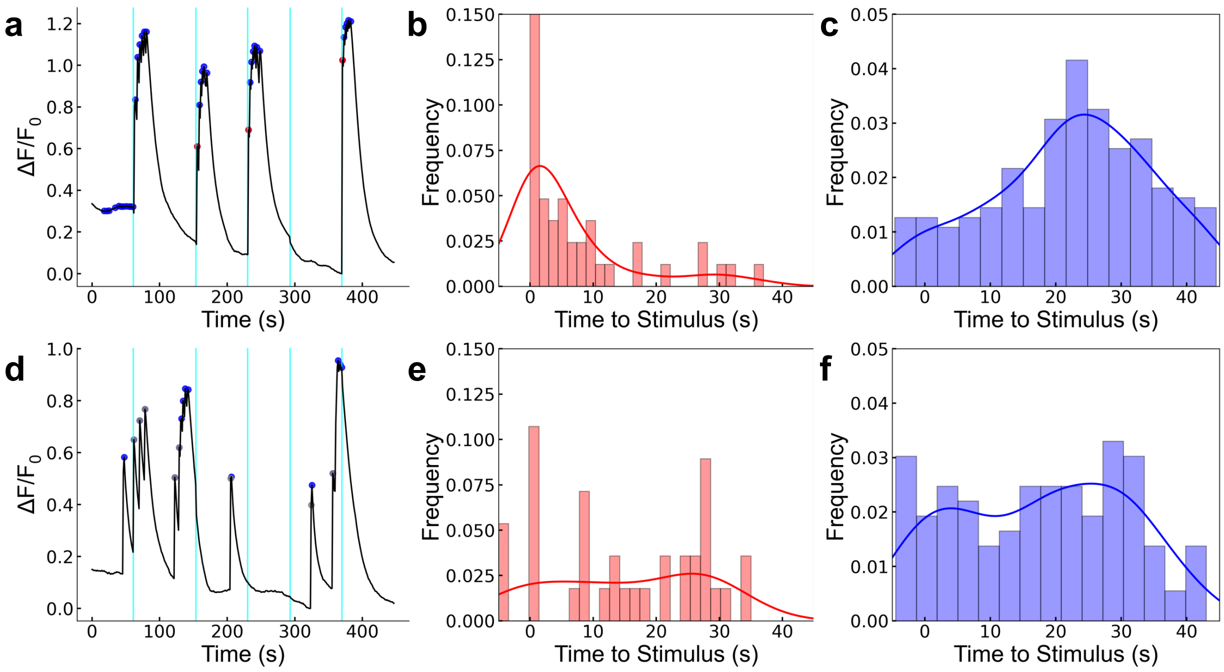


**Supplementary Fig. 21. Peak analysis of MM 4-treated *Hydra* (non-unidirectional MM; rotation rate of 3 MHz).**

**(a)** Characteristic GCaMP7b fluorescence trace of *Hydra* treated with MM **4** + light (Protocol **II**). The contractions (peaks) most often occur in concert with stimuli (cyan lines). **(b)** Temporal location of contraction onset relative to presentation of light stimulus. **(c)** Temporal location of contraction burst peaks relative to presentation of light stimulus. **(d)** Characteristic GCaMP7b fluorescence trace of *Hydra* treated with MM **4** and sham light stimulus (same protocol with 0% laser duty cycle). **(e)** Temporal location of contraction onset peaks relative to presentation of sham light stimulus. Contractions do not occur in concert with sham stimulus. **(f)** Temporal location of contraction burst peaks relative to presentation of sham light stimulus. Histograms were calculated across all collected data (at least 50 stimulation or sham stimulation attempts in at least 5 *Hydra*).

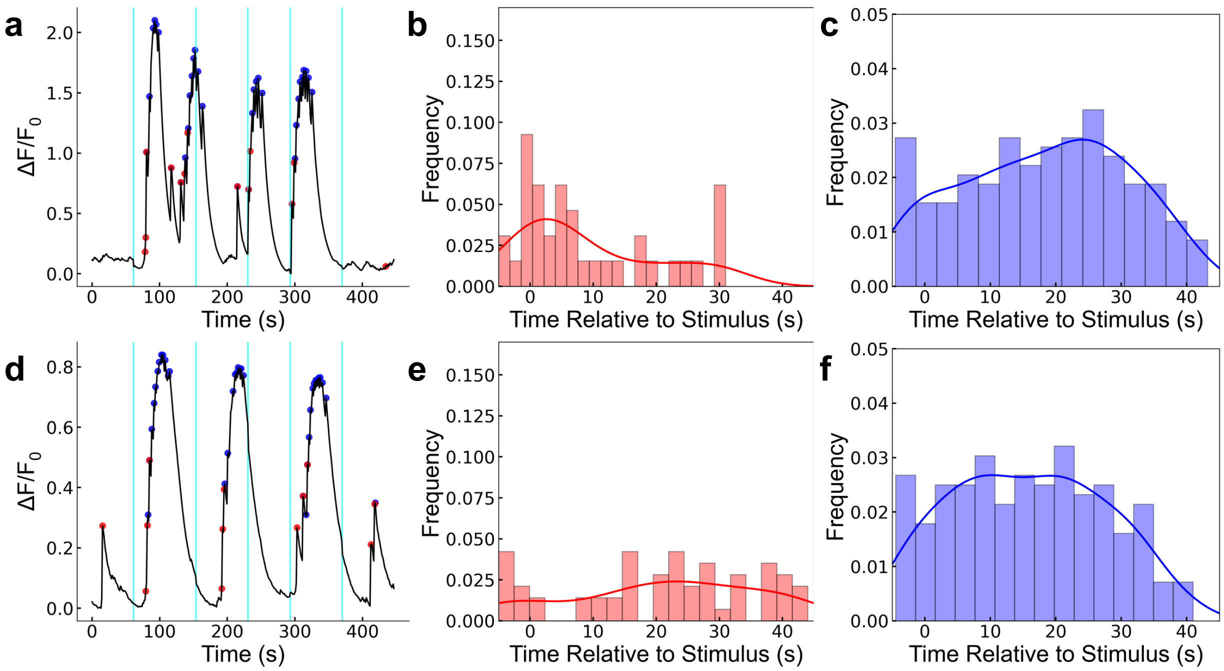


**Supplementary Fig. 22. Peak analysis of DMSO-treated control *Hydra*.**

**(a)** Characteristic GCaMP7b fluorescence trace of *Hydra* treated with DMSO (0.3% v/v) + light (Protocol **II**). The contractions (peaks) occasionally follow stimuli (cyan lines) due to the photosensitivity of the oral region. **(b)** Temporal location of contraction onset relative to presentation of light stimulus. **(c)** Temporal location of contraction burst peaks relative to presentation of light stimulus. **(d)** Characteristic GCaMP7b fluorescence trace of *Hydra* treated with DMSO and sham light stimulus (same protocol with 0% laser duty cycle). Contractions do not occur in concert with sham stimulus. **(e)** Temporal location of contraction onset peaks relative to presentation of sham light stimulus. **(f)** Temporal location of contraction burst peaks relative to presentation of sham light stimulus. Histograms were calculated across all collected data (at least 50 stimulation or sham stimulation attempts in at least 5 *Hydra*).

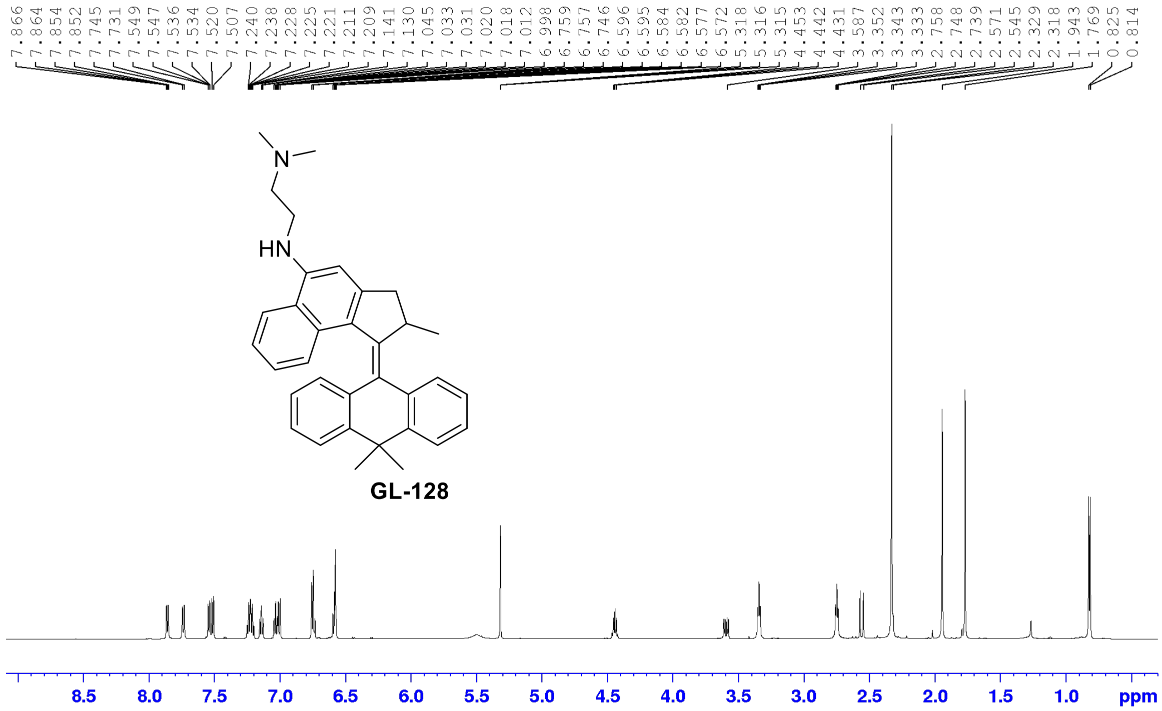


**Supplementary Fig. 23. $^{1}$H NMR spectrum of GL-128 (MM 2).**

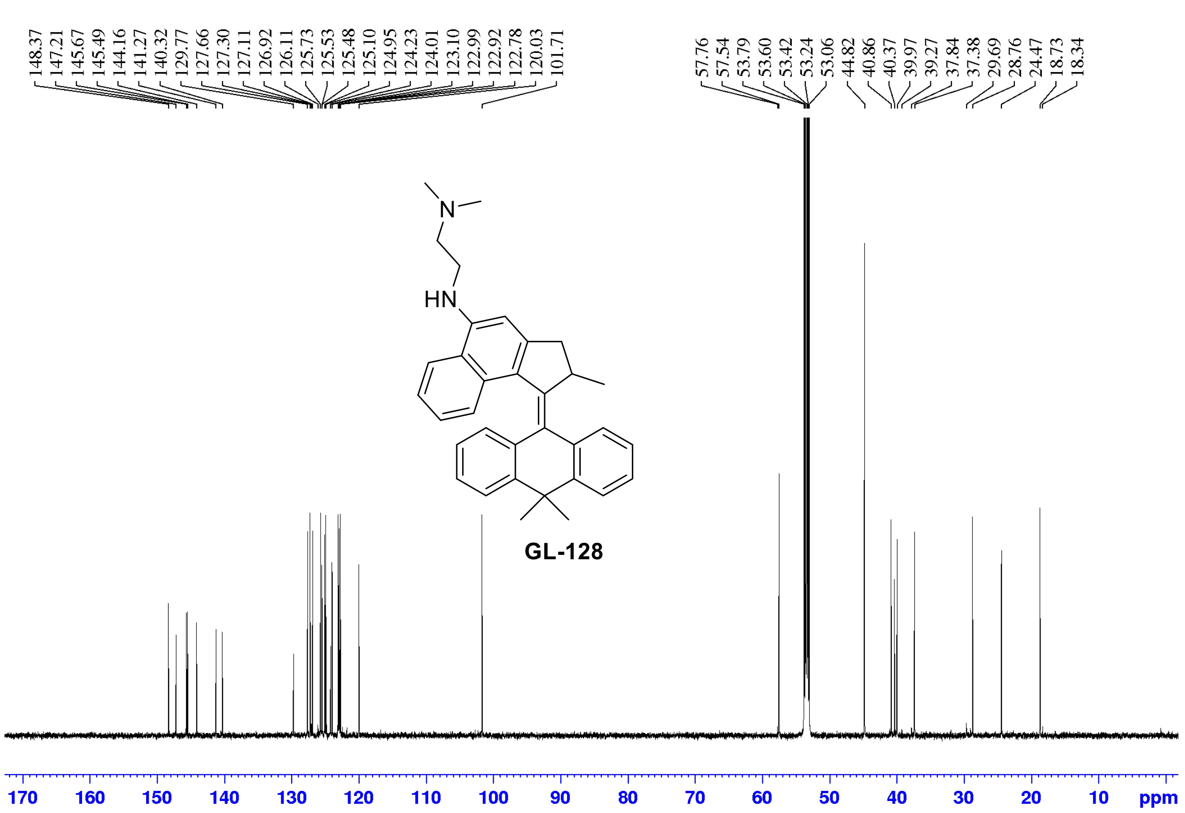


**Supplementary Fig. 24. $^{13}$C NMR of GL-128 (MM 2).**

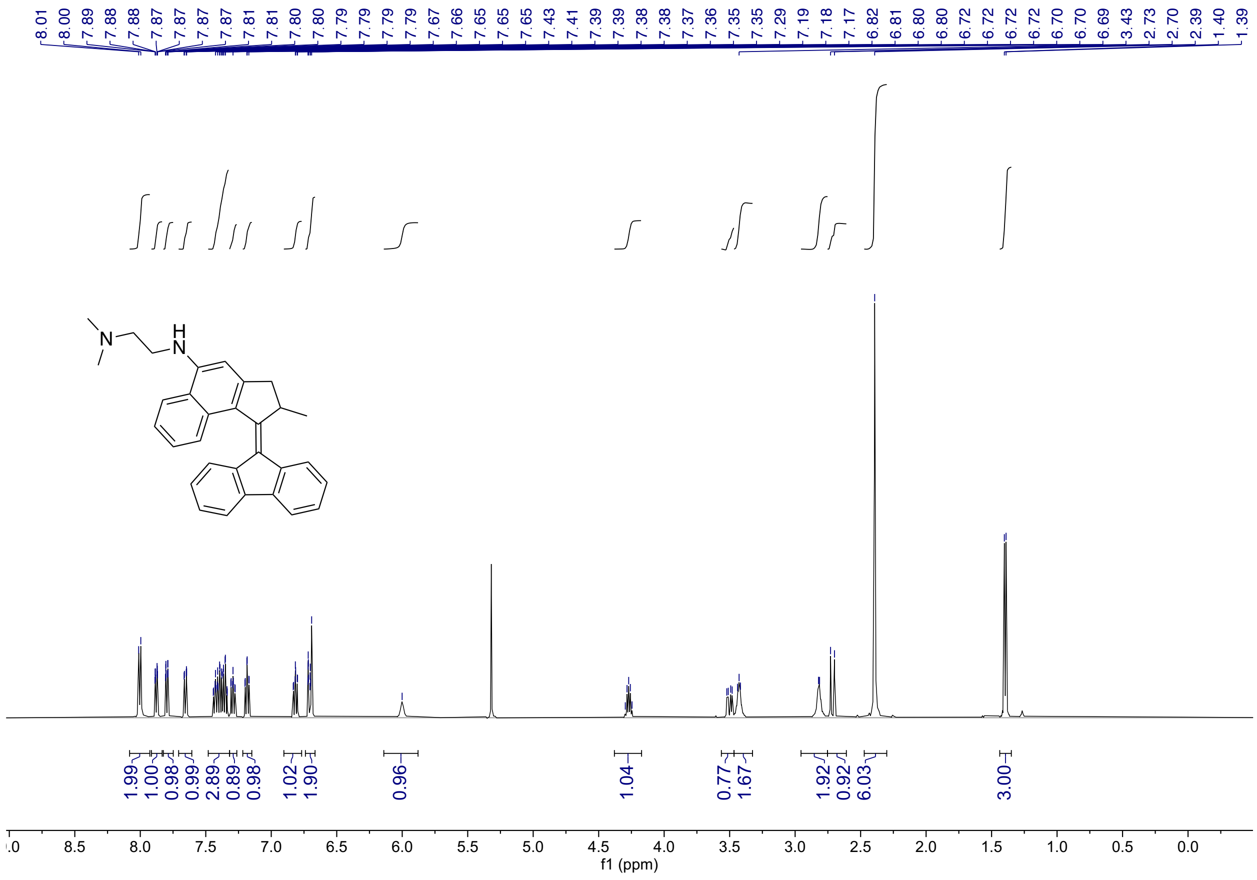

**Supplementary Fig. 25. $^{1}$H-NMR spectrum of MM 3 (ARV-1) in $CD_2Cl_2$**

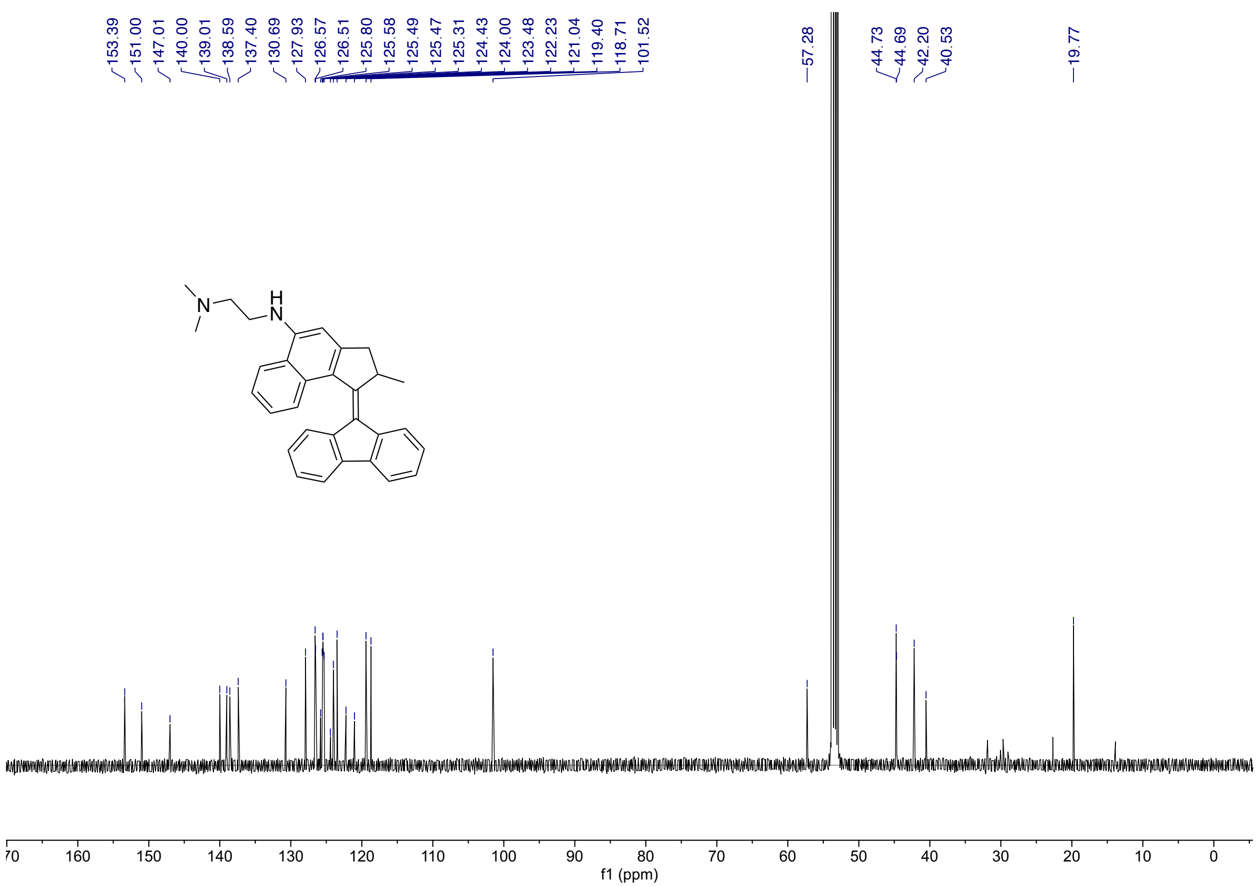


**Supplementary Fig. 26. $^{13}$C-NMR spectrum of MM 3 (ARV-1) in $CD_2Cl_2$**

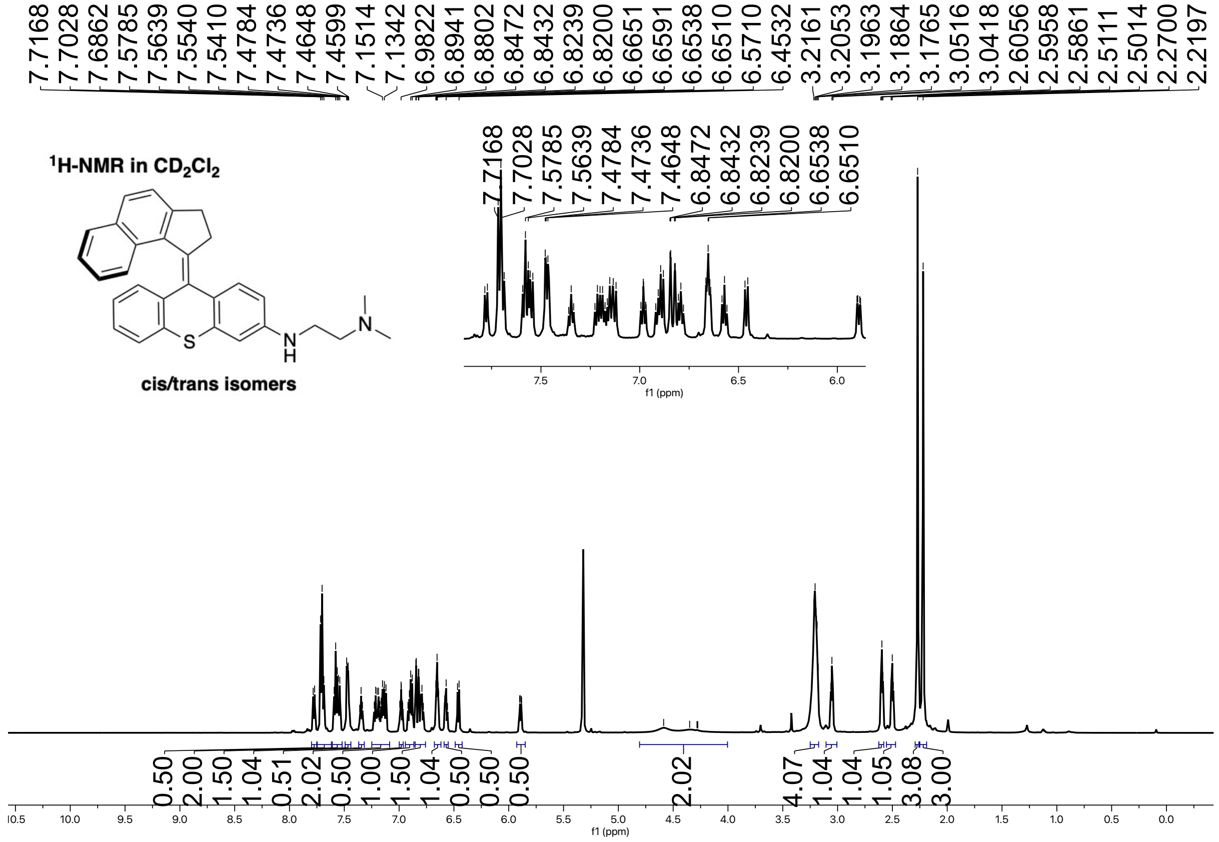


**Supplementary Fig. 27. $^{1}$H-NMR spectrum of MM 4 (BL-167) in $CD_2Cl_2$**

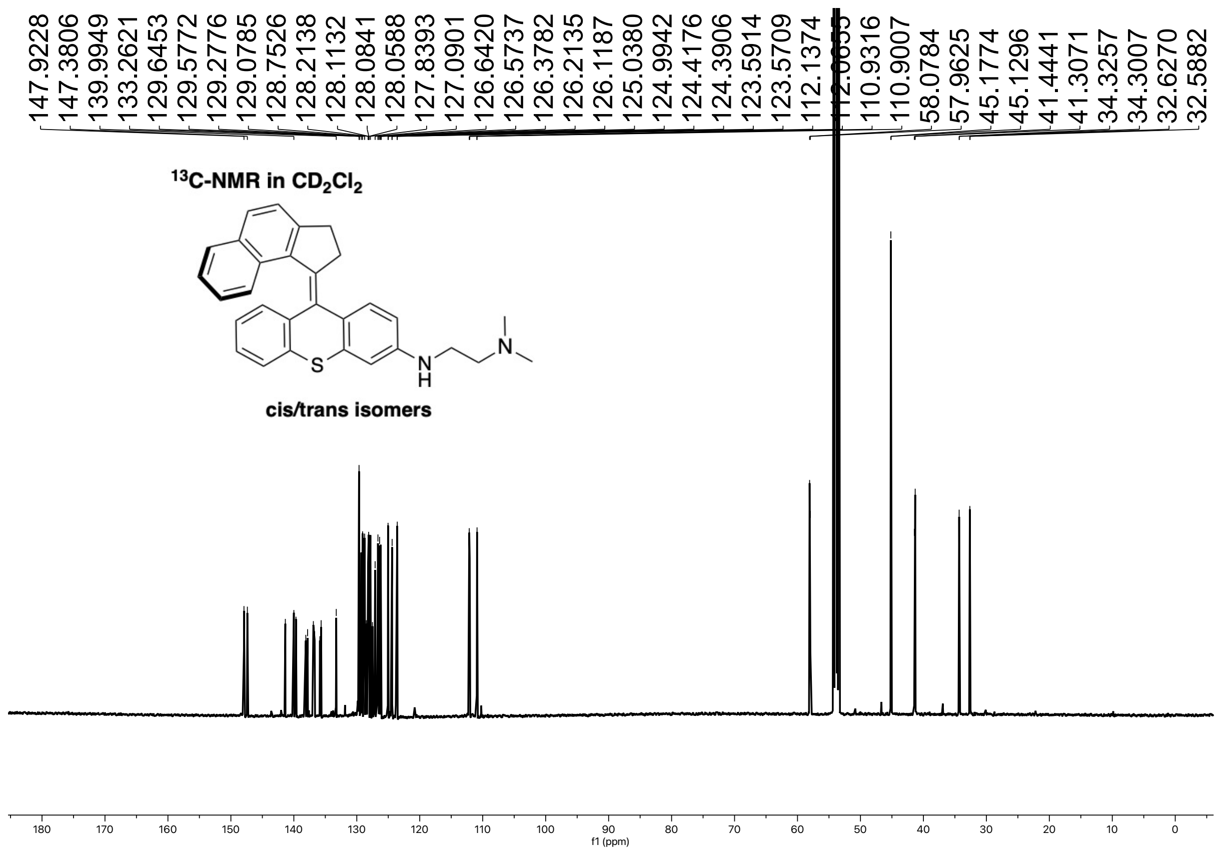


**Supplementary Fig. 28. $^{13}$C-NMR spectrum of MM 4 (BL-167) in $CD_2Cl_2$**

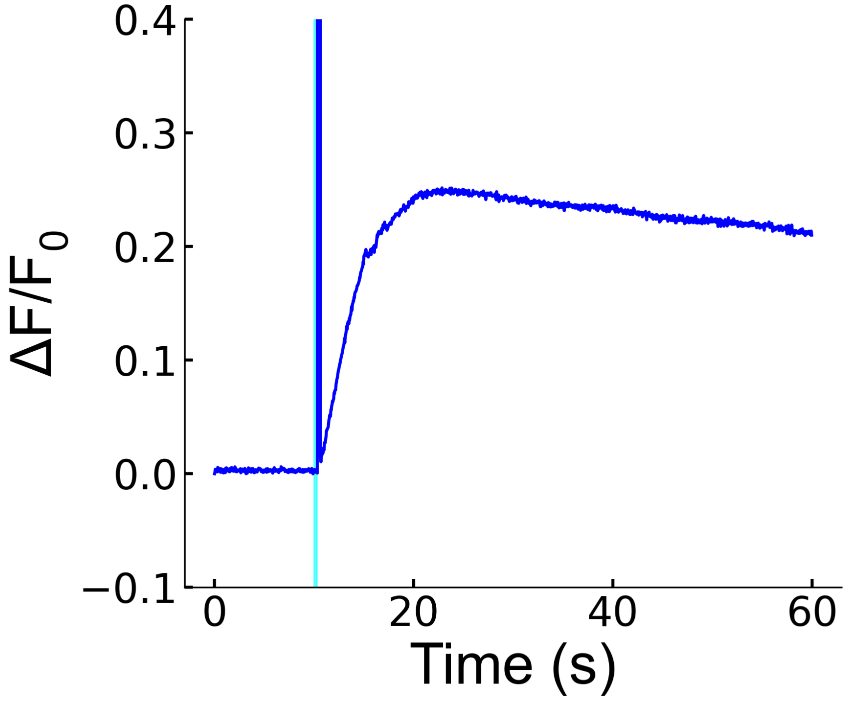


**Supplementary Fig. 29. ICW from a non-scanning laser.**

Normalized fluorescent intensity trace of a HEK293 cell treated with MM **1** as described in the Methods section and excited by a Coherent Chameleon Discovery femtosecond laser operating at ~$1.5\times10^{2}$ W $cm^{-2}$ at 405 nm and imaged at 20 fps. The cyan line indicates the time of stimulation. The sharp vertical line likely reflects a fluorescence resonant energy transfer effect between the MM and calcium imaging dye (Fluo-4) during laser excitation, not visible in other experiments because images were not collected during stimulation.

**Supplementary Table 1. Properties of MM 1-4.**

Partition coefficients and total polar surface area were calculated using the molInspiration online calculator. milogP represents the logarithm of the partition coefficient between octanol and water. Rotation rates were provided using the thermodynamics of the thermal helix inversion step. MM structures provided for reference.

| Molecule | Rotation Rate (Hz) | milogP (a.u.) | Total Polar Surface Area (Å$^2$) | Extinction Coeff @ 400 nm (L mol$^{-1}$cm$^{-1}$) |
|---|---|---|---|---|
| **MM 1** | 3 $\times 10^6$ | 7.90 | 15.27 | ~11,000 |
| **MM 2** | 43 $\times 10^6$ | 7.34 | 15.27 | ~17,500 |
| **MM 3** | 0.1 | 7.31 | 15.27 | ~7,500 |
| **MM 4** | 3 $\times 10^6$ | 7.90 | 15.27 | ~13,000 |

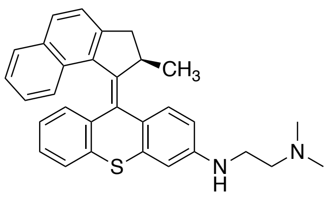


MM **1** (~3 MHz)
Unidirectional

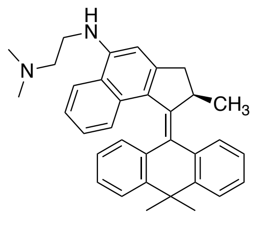


MM **2** (~43 MHz)
Unidirectional

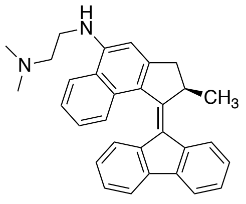


MM **3** (~0.1 Hz)
Unidirectional

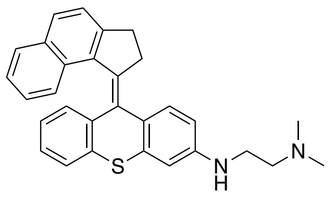


MM **4** (~3 MHz)
Bidirectional

**Supplementary Table 2. Pharmacological blockers and treatments employed.**

| Treatment | Concentration | Process Affected | Target | Process Notes |
|---|---|---|---|---|
| **$Gd^{3+}$** | 50 µM | Extracellular calcium entry | Mechanosensitive Piezo1, Pieoz2, TRPC4 | Dissolved in iECB |
| **Ruthenium Red (RR)** | 10 µM | Extracellular calcium entry | Thermosensitive TRPV1-4 | Dissolved in iECB |
| **PBS** | N/A | Extracellular calcium entry | Extracellular calcium | Replaced iECB |
| **Thapsigargin (Th)** | 1 µM | Intracellular calcium release | Sarco/endoplasmic reticulum pump (SERCA) | Administered in complete growth medium for 1 h prior to exp. |
| **Ryanodine (Ry)** | 100 µM | Intracellular calcium release | Ryanodine receptors (RyR) | Administered in iECB |
| **Xestospongin C (XeC)** | 20 µM | Intracellular calcium release | IP3 receptors (IP3R) | Administered in iECB |
| **U-73122** | 10 µM | Intracellular calcium release | Phospholipase C (PLC) | Administered in iECB |
| **Cytochalasin D** | 2 µM | F-action polymerization | Disrupts PLC->$IP_3R$ spacing and communication | Administered in complete growth |

| | | | | |
|---|---|---|---|---|
| | | | | medium for 2 h prior to exp. |

**Supplementary Table 3.** ***Hydra*** **dataset characteristics.**

| Group | Total Traces | Total *Hydra* Employed | Total Stim. Attempts | Total Successes* | Total Null** | Success Rate*** |
|---|---|---|---|---|---|---|
| **MM 1 + light** | 17 | 10 | 85 | 48 | 14 | 68% |
| **MM 1** | 15 | 5 | 75 | 13 | 28 | 29% |
| **MM 2 + light** | 17 | 10 | 85 | 60 | 16 | 87% |
| **MM 2** | 13 | 6 | 65 | 8 | 25 | 20% |
| **MM 3 + light** | 10 | 5 | 50 | 18 | 2 | 38% |
| **MM 3** | 13 | 8 | 65 | 12 | 17 | 12% |
| **MM 4 + light** | 12 | 6 | 60 | 36 | 5 | 65% |
| **MM 4** | 8 | 5 | 45 | 6 | 15 | 20% |
| **DMSO + light** | 12 | 7 | 60 | 20 | 15 | 44% |
| **DMSO** | 15 | 6 | 75 | 13 | 23 | 21% |

***Successes** were defined as presentations of stimulus that were paired with a contraction and whole-body calcium spike at most three frames (~3 s) after the stimulus was presented. Contractions were identified using a threshold for the change in body GCaMP7b fluorescence ($\Delta F/F_0$ > 1.5x that of the previous data point).

****Null** stimulation attempts were attempts when the stimulus was presented during a period in which the *Hydra* was already contracting ($\Delta F/F_0$ > 0.3).

*****Success rate** was defined as the proportion of stimulation attempts which resulted in a "success", with null attempts discarded.

For more info see *Methods*.

**Supplementary Movie 1.**

Timelapse of Fluo-4 fluorescence in cardiomyocytes treated with MM **1** (8 μM) and Fluo-4 and stimulated with 400 nm laser light for 250 ms at an irradiance of $5.1\times10^{2}$ W $cm^{-2}$. The cell on the left is stimulated at 30 s.

**Supplementary Movie 2.**

Timelapse of bright field images of cardiomyocytes treated with MM **1** (8 μM) and Fluo-4 and stimulated with 400 nm laser light for 250 ms at an irradiance of $5.1\times10^{2}$ W $cm^{-2}$. The cell on the left is stimulated at 30 s.

**Supplementary Movie 3.**

A *Hydra* expressing GCaMP7b in epitheliomuscular cells treated with MM **1** (24 μM) and stimulated with 405 nm laser light for 1 s at an irradiance of $9.0\times10^{2}$ W $cm^{-2}$ (Protocol **I**). Stimulation is applied at 10 s. The blue circle shown at 10 s corresponds to the area of stimulation. The trace shown at the bottom of the video corresponds to the intensity of GCaMP7b fluorescence in the area of stimulation across the timescale of the video. The cyan line in the trace represents the time of stimulus presentation, during which no data was collected.

**Supplementary Movie 4.**

A *Hydra* expressing GCaMP7b in epitheliomuscular cells treated with DMSO solvent vehicle (0.3% v/v) and stimulated with 405 nm laser light for 1 s at an irradiance of $9.0\times10^{2}$ W $cm^{-2}$ (Protocol **I**). Stimulation is applied at 10 s. The blue circle shown at 10 s corresponds to the area of stimulation. The trace shown at the bottom corresponds to the intensity of GCaMP7b fluorescence in the area of stimulation across the timescale of the video. The cyan line in the trace represents the time of stimulus presentation, during which no data was collected.

**Supplementary Movie 5.**

A *Hydra* expressing GCaMP7b in epitheliomuscular cells treated with MM **2** (24 µM) and stimulated with 405 nm laser light for 1 s at an irradiance of $6.8 \times 10^2$ W $cm^{-2}$ (Protocol **I**). Stimulation is applied at 10 s at two distinct locations in the *Hydra* indicated by the yellow and blue circles that appear during stimulation. The traces shown at the bottom of the video correspond to the intensity of GCaMP7b fluorescence in the areas of stimulation across the timescale of the video.

**Supplementary Movie 6.**

A *Hydra* expressing GCaMP7b in epitheliomuscular cells treated with MM **2** (24 µM) and stimulated with 405 nm laser light for 2 s at an irradiance of $9.0 \times 10^2$ W $cm^{-2}$. Light was delivered to the oral region of the *Hydra* (Protocol **II**) at irregular intervals. The yellow circle around the oral region indicates the region to which stimulation was applied. The trace shown at the bottom of the video corresponds to the intensity of GCaMP7b fluorescence in the whole *Hydra* across the timescale of the video. The vertical yellow lines in the trace mark times during which stimulus was presented, during which no data was collected. Contractile and electrophysiological responses were observed upon each presentation of stimulus.
